# Insulin receptor substrate 2 (IRS2) confers resistance to PI3K pathway inhibition in *PIK3CA* mutant breast cancer

**DOI:** 10.64898/2026.04.24.720709

**Authors:** Michael W. Lero, Jennifer S. Morgan, Michael-Anthony Card, Lihua Julie Zhu, Junhui Li, Rui Li, Quyen Thu Bui, Emma Mohlmann, Leslie M. Shaw

## Abstract

Activating mutations in PI3K are one of the most frequent mutations in breast cancer and are associated with worse patient outcomes in many breast cancer subtypes. Despite intense interest, cancer treatments that target the PI3K pathway have been only modestly effective due to intrinsic and acquired resistance mechanisms which reactivate PI3K signaling. Here, we characterize a feedback mechanism by which PI3K pathway inhibitors increase insulin receptor substrate 2 (IRS2) abundance and demonstrate the role of IRS2 in promoting resistance to these drugs. In *PIK3CA* mutant breast tumors and cell lines, there is a significant reduction in IRS2 mRNA and protein abundance which is reversed by PI3K pathway inhibition and mediated by the transcription factor FOXO3. *PIK3CA* mutations do not alter IRS1 expression. IRS2 confers resistance to PI3K pathway inhibition by sustaining PI3K signaling in *PIK3CA* mutant, but not wild-type breast cancer cells. Increased IRS2 abundance also correlates with PI3K pathway inhibitor resistance across PI3K mutant cancer cell lines from a variety of tissues. The clinical relevance of these findings is highlighted by the frequency of PI3K mutations in cancer and the identification of a new target to address the challenges associated with prior efforts to block the reactivation of PI3K signaling during PI3K inhibition.

## Introduction

There continues to be intense interest in understanding how phosphatidylinositol-3 kinase (PI3K) contributes to tumorigenesis and how it can be exploited therapeutically. *PIK3CA*, which encodes the p110α catalytic subunit of PI3K, is frequently mutated in human cancer, and is the second most commonly mutated gene in breast cancer(1). Three hotspot mutations, E545K, H1047R, and H1047L, account for 56% of all PI3K mutations in breast cancer(2). These activating mutations cause constitutive PI3K signaling, leading to growth-factor independent cell proliferation, increased cell survival and invasion, and generally worse clinical outcomes compared to patients with *PIK3CA* wild-type tumors(3–13). Two PI3K inhibitors have been shown to extend progression free survival in select *PIK3CA* mutant breast cancers and have received FDA approval(14–17). However, acquired and intrinsic resistance mechanisms that sustain PI3K signaling during treatment significantly limit the clinical efficacy of these drugs.

Many clinically observed mechanisms of acquired resistance to PI3K inhibitors cause reactivation of PI3K signaling, either through secondary *PIK3CA* mutations, *AKT1* activating mutations, *PTEN* deletions, or *PTEN* loss of function mutations(18–20). Even in the absence of genomic alterations, PI3K pathway inhibitors elicit rapid intrinsic resistance mechanisms that reactivate PI3K signaling and limit drug efficacy. Because of the role of PI3K in glucose uptake and metabolism, PI3K inhibitors raise blood glucose levels, which leads to increased pancreatic insulin production and elevated blood insulin levels(21, 22). This excess blood insulin has been implicated in signaling through the insulin receptor to reactivate the PI3K pathway and promote tumor growth during PI3K inhibition in murine models of cancer(21, 23). PI3K pathway inhibitors also relieve negative feedback mechanisms that would otherwise limit PI3K signaling. This leads to many changes, including upregulation of the insulin-like growth factor 1 receptor (IGF1R), which reactivates the PI3K pathway. Interruption of this upstream IGF1R signaling has been shown to sensitize cancer to PI3K and AKT inhibition in vivo(24–32). Despite the fact that clinical PI3K-IGF1R treatment combinations have been discontinued because of concerns with side effects, these observations highlight the importance of both receptor-dependent and downstream reactivation of PI3K signaling in mediating resistance to PI3K inhibition(33).

This study was motivated by our interest in the potential contribution of the insulin receptor substrate (IRS) proteins to resistance to PI3K inhibition. The insulin and IGF1 receptors signal through IRS1 and IRS2 to activate PI3K/AKT signaling(34–36). Importantly, while both IRS1 and IRS2 activate PI3K and facilitate glucose homeostasis, IRS2 uniquely promotes breast cancer cell invasion and cancer stem cell self-renewal through ligand-dependent activation of PI3K signaling(37, 38). These functional differences raise the possibility that IRS1 and IRS2 may also have divergent contributions to PI3K pathway inhibitor sensitivity. Because of the importance of upstream PI3K signaling in mediating resistance to PI3K inhibition, and the role of IRS proteins in receptor-mediated PI3K signaling, we investigated the interplay among *PIK3CA* mutations, PI3K pathway inhibitors, and IRS proteins in breast cancer. We report that IRS2 expression confers resistance to PI3K pathway inhibition in *PIK3CA* mutant, but not *PIK3CA* WT breast cancer.

## Results

### PI3K signaling controls IRS2 mRNA and protein abundance in *PIK3CA* mutant breast cancer

To assess the impact of activating *PIK3CA* mutations on IRS adaptor proteins, we first examined whether *PIK3CA* mutations were associated with IRS1 or IRS2 abundance in breast cancer patient samples from the METABRIC dataset. Whereas *IRS1* mRNA levels were similar in WT and mutant *PIK3CA* tumors, mutations in *PIK3CA* correlated with significant reductions in *IRS2* mRNA in luminal breast cancer, triple-negative breast cancer (TNBC), and HER2-enriched breast cancer (Fig 1A,B, Fig S1A)(39), findings consistent with previous reports(40, 41). This phenotype was also present in the independent CPTAC dataset, where *PIK3CA* mutations in breast cancer correlated significantly with reductions in both IRS2 mRNA and protein (Fig 1C,D)(42). Additional genomic alterations that lead to PI3K/AKT pathway activation, including *PTEN* deletion and *PTEN* mutation, also correlated with reduced *IRS2* mRNA in breast cancer patient samples (Fig S1B-E)(39).

**Figure 1.**
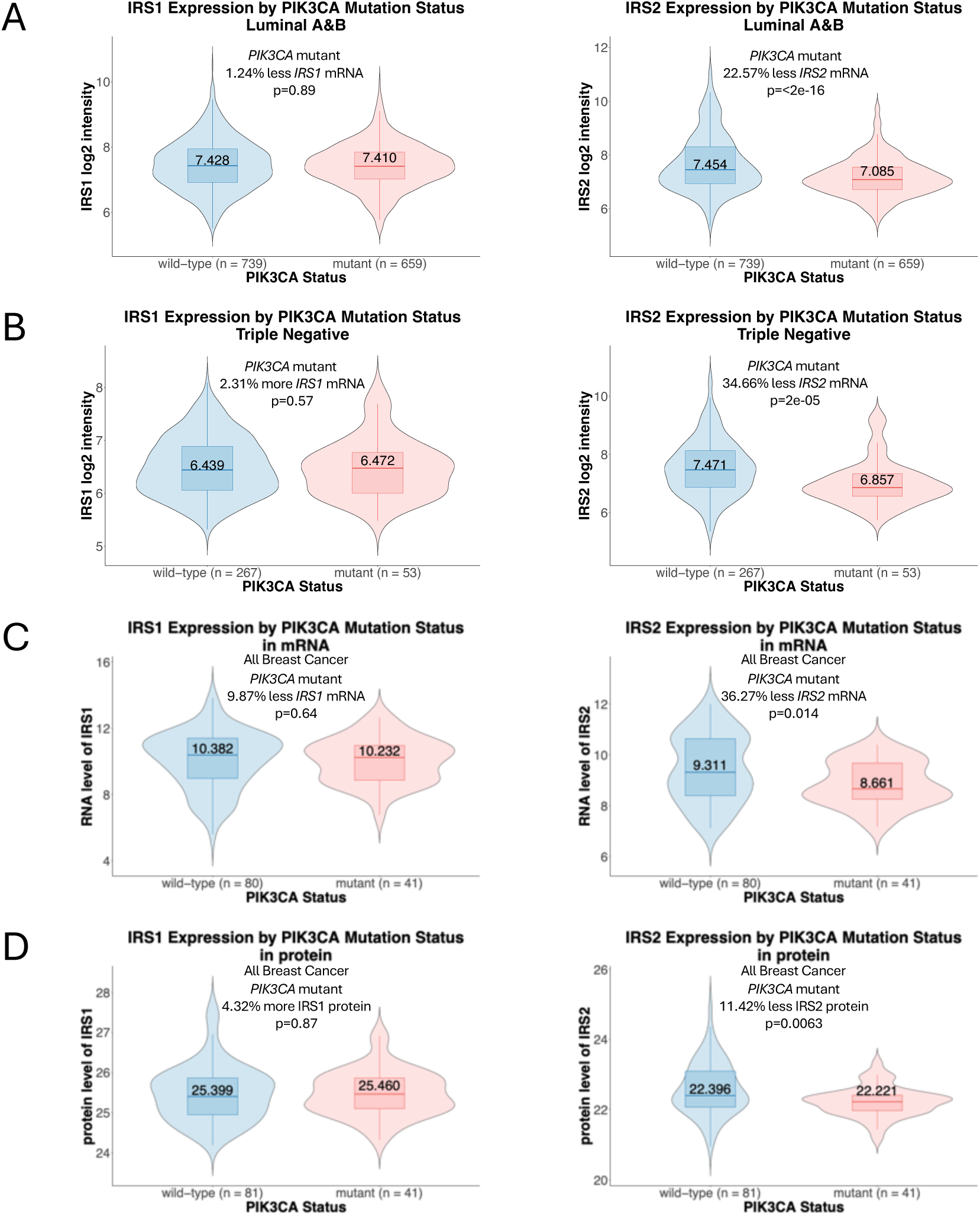
Reduction of IRS2 mRNA and protein abundance in *PIK3CA* mutant breast cancer. Differential gene expression analysis of IRS1 and IRS2 in *PIK3CA* WT and *PIK3CA* mutant luminal A&B (A) and triple-negative (B) breast tumors in the METABRIC dataset. Gene expression measurements shown as log2 intensity values (Illumina HT-12 v3 microarray). Differential mRNA (C) and protein (D) expression analysis of IRS1 and IRS2 in *PIK3CA* WT and *PIK3CA* mutant breast tumors in the Clinical Proteomic Tumor Analysis Consortium (CPTAC) dataset (comparisons include all breast cancer types). mRNA expression shown as Log2(RSEM+1), protein expression shown as Log2(MS1 intensity). For all violin plots, the box and whisker plots show the median, 25%(Q1), 75%(Q3) of data (box), Q1 - 1.5 × interquartile range (IQR), and Q3 + 1.5 × IQR of data (whiskers). For all panels, percent changes were determined by comparing the medians of the original data (prior to log transformation). For all panels, statistical analysis by Wilcoxon rank sum test with continuity correction. See also Figure S1.

Tumors often harbor multiple oncogenic mutations which make it difficult to attribute a given observation to a specific genomic alteration. To determine the isolated impact of activating *PIK3CA* mutations on IRS1 and IRS2 abundance, we utilized non-tumorigenic MCF10A human mammary epithelial cells expressing WT *PIK3CA* or heterozygous knock-in of the two most common *PIK3CA* activating mutations, E545K or H1047R(2, 4). Importantly, these cells are isogenic, ensuring that any differential phenotypes are due to *PIK3CA* mutational status alone and not to other differences in mutational background. Two single-cell clones of each knock-in mutant were studied to control for clonal differences. *PIK3CA* mutant cells exhibited elevated PI3K/AKT signaling (Fig S2A). Wild-type and *PIK3CA* mutant cells expressed equivalent levels of IRS1 mRNA and protein in three of the four mutant clones, with one *PIK3CA* mutant clone expressing elevated IRS1 (Fig 2A,B). In contrast, all *PIK3CA* mutant clones expressed significantly lower IRS2 mRNA and protein levels compared to wild-type MCF10A cells (Fig 2A,B).

**Figure 2.**
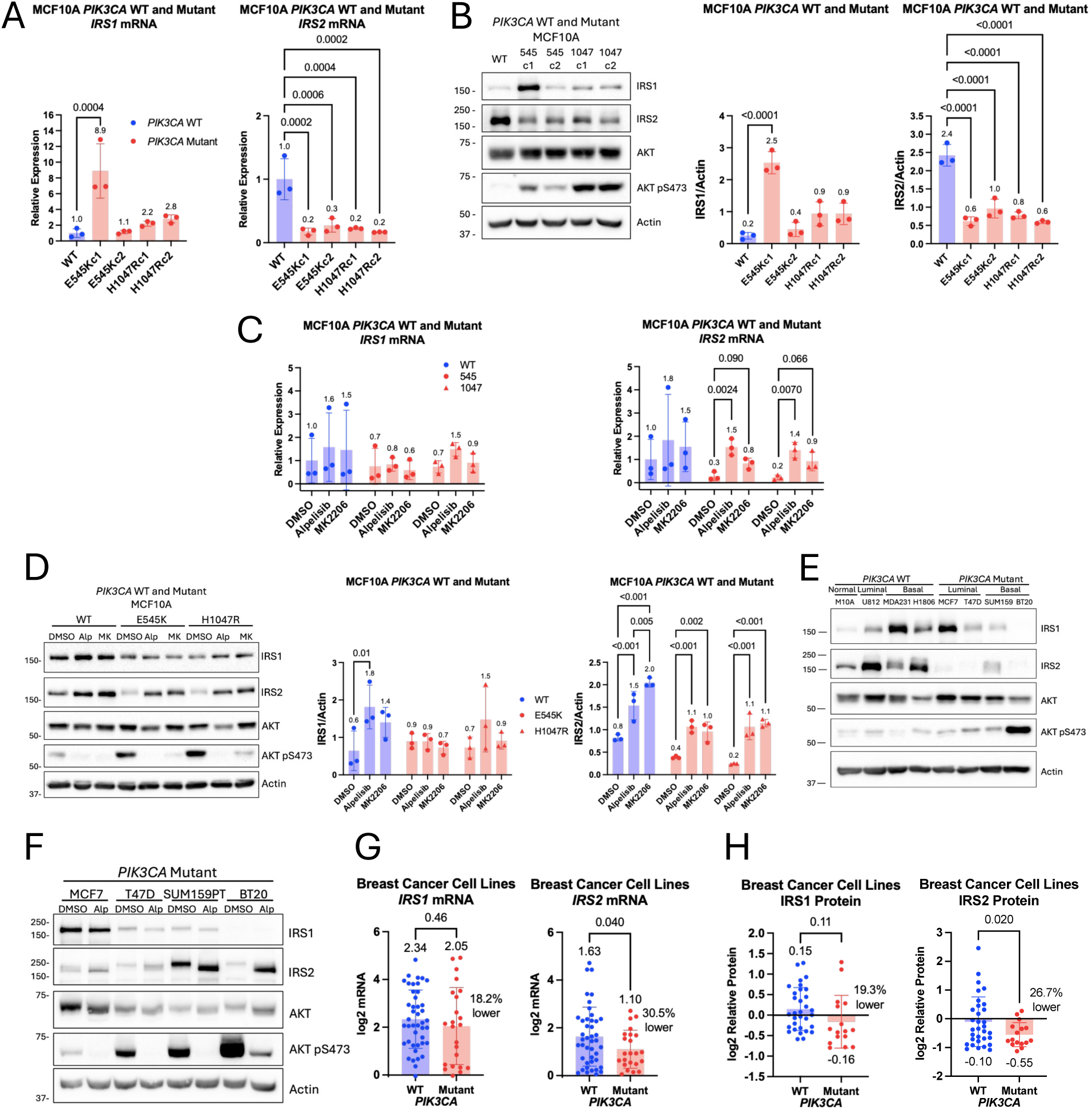
*PIK3CA* mutation reversibly suppresses IRS2 mRNA and protein abundance in mammary epithelial cells and breast cancer cell lines. (A and B) *PIK3CA* WT and *PIK3CA* mutant knockin MCF10A cells were serum starved, then evaluated for IRS1 and IRS2 mRNA (A) and protein (B) expression. The data shown in the graphs represent the mean +/- standard deviation (SD) of three independent experiments. Statistical analysis by One-Way ANOVA followed by Dunnett’s multiple comparisons test to compare mutants against WT (P values shown). (C and D) *PIK3CA* WT and *PIK3CA* mutant knockin MCF10A cells were treated in plain media with or without alpelisib (5μM) or MK2206 (1μM) for 24 hrs and then evaluated for IRS1 or IRS2 mRNA (C) or protein (D) expression. The data shown in the graphs represent the mean +/- SD of three independent experiments. Statistical analysis by (C) separate One-Way ANOVA followed by the Bonferroni test for multiple comparisons to compare drug treatments to DMSO or (D) Two-Way ANOVA followed by the Tukey test for multiple comparisons to compare all treatments within each cell type (P values shown). (E) IRS1 and IRS2 protein expression in *PIK3CA* WT and *PIK3CA* mutant breast epithelial and cancer cell lines following serum starvation (M10A - MCF10A, U812 - UACC812, MDA-231 – MDA-MB-231, H1806 - HCC1806, SUM159 - SUM159PT). (F) *PIK3CA* mutant breast cancer cells were treated in plain media with or without alpelisib (5μM) for 24 hrs and cell extracts were evaluated by immunoblot. (G and H) Differential IRS1 and IRS2 mRNA (G) and protein (H) in breast cancer cell lines from DepMap. (mRNA - log2(TPM+1), protein RPPA signal). The data shown in the graphs represent the mean +/- SD of all included cell lines. Percent changes were determined by comparing the means of the original data (prior to log transformation). Statistical analysis by Welch’s t test (two-tailed) (P values shown). See also Figure S2.

To assess if the reduction in IRS2 expression was related to increased PI3K signaling in *PIK3CA* mutant cells, *PIK3CA* wild-type and mutant cells were treated with a clinically approved PI3Kα specific inhibitor (alpelisib) or an AKT1/2/3 inhibitor (MK2206). While both PI3Kα and AKT inhibition reduced AKT signaling as evidenced by lower AKT pS473 levels, neither PI3Kα nor AKT inhibition altered IRS1 mRNA or protein levels in *PIK3CA* mutant cells (Fig 2C,D and Fig S2B). In contrast, both PI3Kα and AKT inhibition led to increased IRS2 mRNA and protein in wild-type as well as E545K and H1047R *PIK3CA* mutant cells (Fig 2C,D). This is consistent with a prior report of PI3K inhibition leading to increased *IRS2* mRNA in *PIK3CA* mutant cell lines(41).

While the use of MCF10A cells enabled us to assess the contribution of *PIK3CA* mutations to IRS1 and IRS2 abundance in cells lacking other oncogenic genomic alterations, they are not cancer cells. To determine if mutant *PIK3CA* also regulated IRS2 expression in breast cancer cell lines, IRS protein abundance was measured in three *PIK3CA* wild-type breast cancer cell lines and four *PIK3CA* mutant breast cancer cell lines. IRS1 expression was variable across all cell lines but IRS2 expression was markedly reduced in all four *PIK3CA* mutant cell lines compared to wild-type *PIK3CA* cells (Fig 2E). In each *PIK3CA* mutant cell line, PI3Kα inhibition yielded elevated IRS2 protein levels compared to vehicle treated controls (Fig 2F). In contrast, PI3Kα inhibition in *PIK3CA* wild-type cells did not consistently alter IRS2 protein abundance (Fig S2C). To assess if this selective *PIK3CA* phenotype was present in a larger number of breast cancer cell lines, we analyzed publicly available data from the Cancer Dependency Map (DepMap)(43). In these data, *PIK3CA* mutations were also associated with significantly reduced IRS2 mRNA and protein abundance, but IRS1 expression showed no association (Fig 2G,H).

#### Mutant *PIK3CA* alters IRS2 expression through FOXO3

The strong influence of *PIK3CA* mutations and PI3K pathway inhibitors on *IRS2* mRNA abundance led us to hypothesize that active PI3K signaling regulates IRS2 expression through transcription factor regulation. While other IRS2 transcription factors have been described, we focused on FOXO1 and FOXO3 as they are regulated by PI3K/AKT signaling and do not drive IRS1 expression(44, 45). PI3K signaling and *PIK3CA* mutations activate AKT, which phosphorylates FOXO1/3 in the nucleus, leading to FOXO1/3 nuclear exclusion and interrupting the ability of FOXO1/3 to facilitate target gene expression(24, 44–49). To determine if FOXO1/3 contribute to the PI3K signaling-dependent changes in *IRS2* mRNA levels that we observed, the influence of *PIK3CA* mutations on expression and phosphorylation of FOXO1 (pS256) and FOXO3 (pS253) was assessed. In *PIK3CA* mutant breast tumor cell lines, PI3Kα inhibition led to reduced phosphorylation of FOXO1 (S256) and FOXO3 (S253), indicating that mutant *PIK3CA* was responsible for mediating FOXO1/3 inhibitory phosphorylation (Fig 3A). PI3Kα inhibition in *PIK3CA* H1047R MCF10A cells also led to increased nuclear FOXO1 and FOXO3. IRS2 nuclear localization also increased, which we have previously reported to be regulated in a PI3K-dependent manner (Fig 3B)(50).

**Figure 3.**
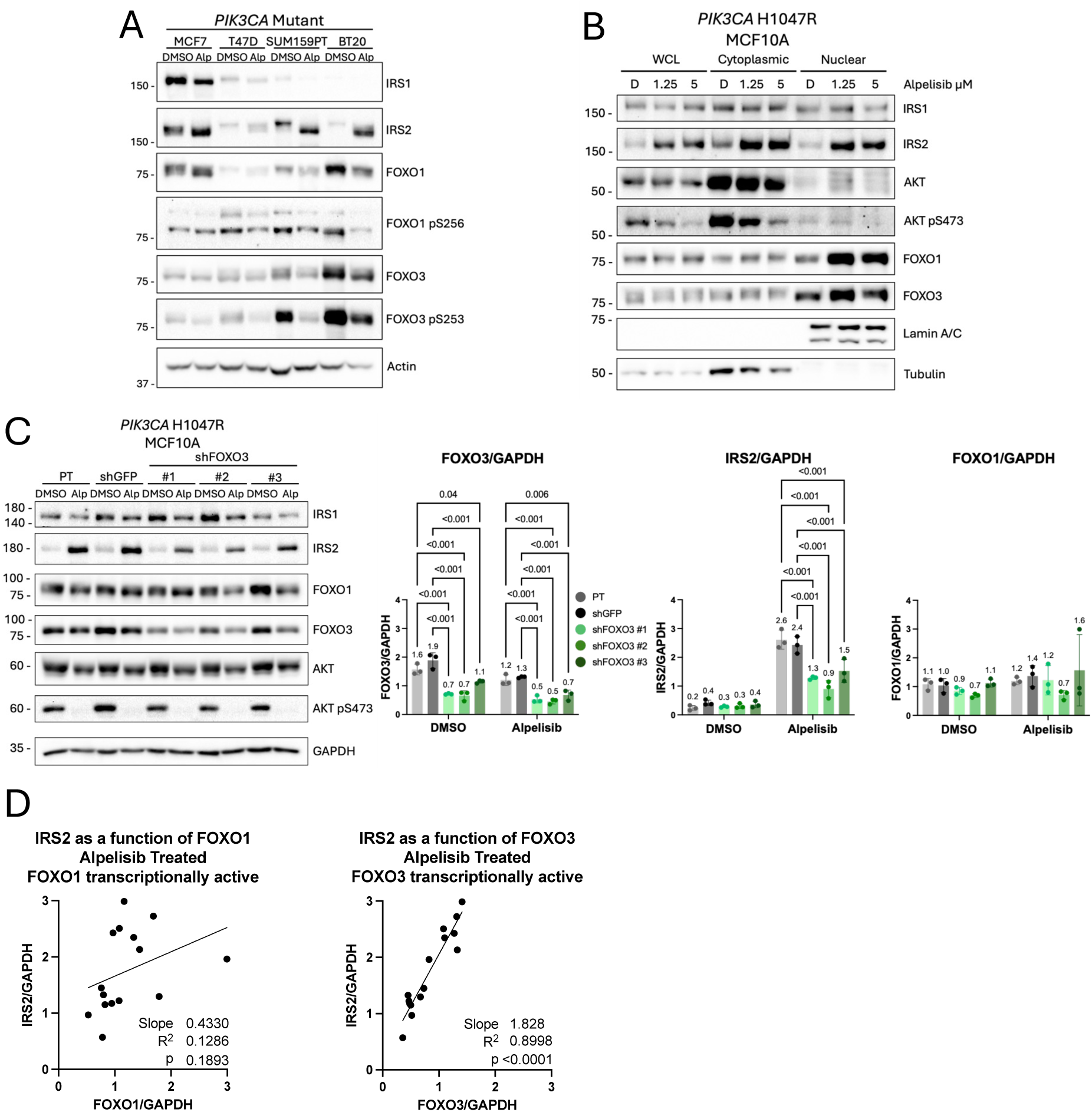
Mutant *PIK3CA* regulates IRS2 expression through FOXO3. (A) *PIK3CA* mutant breast cancer cell lines were treated in plain media with or without alpelisib (5μM) for 24 hrs and cell extracts were evaluated by immunoblot. (B) *PIK3CA* mutant H1047R MCF10A cells were treated in plain media with or without alpelisib for 24 hours and cytoplasmic and nuclear extracts were evaluated by immunoblot (D = DMSO). (C) *PIK3CA* mutant H1047R MCF10A parental (PT) cells as well as cells expressing control shRNA (shGFP) or shRNA targeting FOXO3 were treated in plain media with or without alpelisib (5μM) for 24 hrs and cell extracts were evaluated by immunoblot. The data shown in the graphs represent the mean +/- SD of three independent experiments. Statistical analysis by Two-Way ANOVA followed by the Bonferroni test for multiple comparisons, comparing each control (PT and shGFP) to all other sublines within each treatment (P values shown). (D) Association of IRS2 expression with FOXO1 and FOXO3 abundance in alpelisib treated cells from (C). Statistical analysis by simple linear regression, p<0.05 indicates that the slope is significantly different from zero. See also Figure S3.

To further dissect the mechanism of increased *IRS2* mRNA levels in response to PI3K inhibition we focused on FOXO3, based upon previous reports implicating this transcription factor as playing a predominant role in the feedback regulation of IRS2 expression(44, 51–54). FOXO3 knock-down was achieved using three independent shRNAs (Fig 3C). Levels of IRS2 did not differ between vehicle-treated control and FOXO3 knock-down (KD) cells, suggesting that FOXO3 is not responsible for basal IRS2 expression in *PIK3CA* mutant cells. However, the increase in IRS2 expression observed in response to PI3Kα inhibition was attenuated in FOXO3 KD cells. Importantly, FOXO1 expression levels were not altered in FOXO3 KD cells. Expression of IRS1 was also not changed (Fig S3A).

Linear regression was performed to assess the relationship between IRS2 expression and FOXO1 or FOXO3 abundance. In this analysis, normalized IRS2 protein level was plotted against the normalized FOXO1 or FOXO3 protein abundance for all replicates of all control and FOXO3 KD cells (each dot is from one replicate of control or FOXO3 KD cells). Because PI3K signaling renders FOXO1/3 transcriptionally inactive, vehicle treated and alpelisib treated samples were plotted separately to interrogate FOXO1/3-dependent IRS2 expression in different signaling contexts. When only vehicle treated samples were plotted, FOXO1 and FOXO3 expression was not correlated with IRS2 abundance, indicating that these transcription factors do not drive IRS2 expression at baseline in *PIK3CA* mutant cells (Fig S3B). When PI3K inhibitor-treated samples were plotted, FOXO1 levels were weakly correlated with IRS2 abundance (Fig 3D). In contrast, FOXO3 levels showed a high correlation with IRS2 abundance, indicating that over 89% of the variation in IRS2 was explained by FOXO3 (R^2^=0.8998) (Fig 3D). Together, these results suggest that in a *PIK3CA* mutant context, while PI3K signaling affects transcriptional activity and localization of both FOXO1 and FOXO3, FOXO3 is primarily responsible for PI3K signaling-dependent changes in *IRS2* mRNA levels, similar to what has been observed in pancreatic β-cells(44).

### IRS2 confers resistance to PI3K pathway inhibitors in *PIK3CA* mutant breast cancer

IRS2 is an adaptor protein that links upstream receptor activation with PI3K/AKT signaling. While *PIK3CA* mutations increase baseline PI3K signaling, ligand stimulation can further augment PI3K activation, which is important given the role of sustained PI3K signaling in resistance to PI3K inhibitors (Fig S4A,B). To assess the role of IRS2 in resistance to PI3K pathway inhibition, we first generated control (NT) and IRS2-null (IRS2KO) MDA-MB-231 (*PIK3CA* WT) and SUM159PT (*PIK3CA* H1047L) cell lines using CRISPR-Cas9 gene editing (Fig 4A,B). Cells were plated, treated with increasing doses of PI3K pathway inhibitors for 72 hours, fixed, and stained with crystal violet to measure the size of the cell population following drug treatment. Half-maximal inhibitory concentrations (IC50s) were estimated and used to compare drug sensitivity in different experimental conditions. Variable slope nonlinear regression was used to determine a best fit curve for the data and to estimate IC50 values(55). When *PIK3CA* wild-type cells were treated with alpelisib, control and IRS2KO cells exhibited equivalent drug sensitivity (Fig 4C). This was also true when *PIK3CA* wild-type cells were treated with the AKT1/2/3 inhibitor MK2206 (Fig 4C). In marked contrast, *PIK3CA* mutant (H1047L) cells lacking IRS2 exhibited increased sensitivity to PI3K and AKT inhibition compared to control cells, with IRS2KO IC50 values less than half of control IC50 values for alpelisib and MK2206 (Fig 4D).

**Figure 4.**
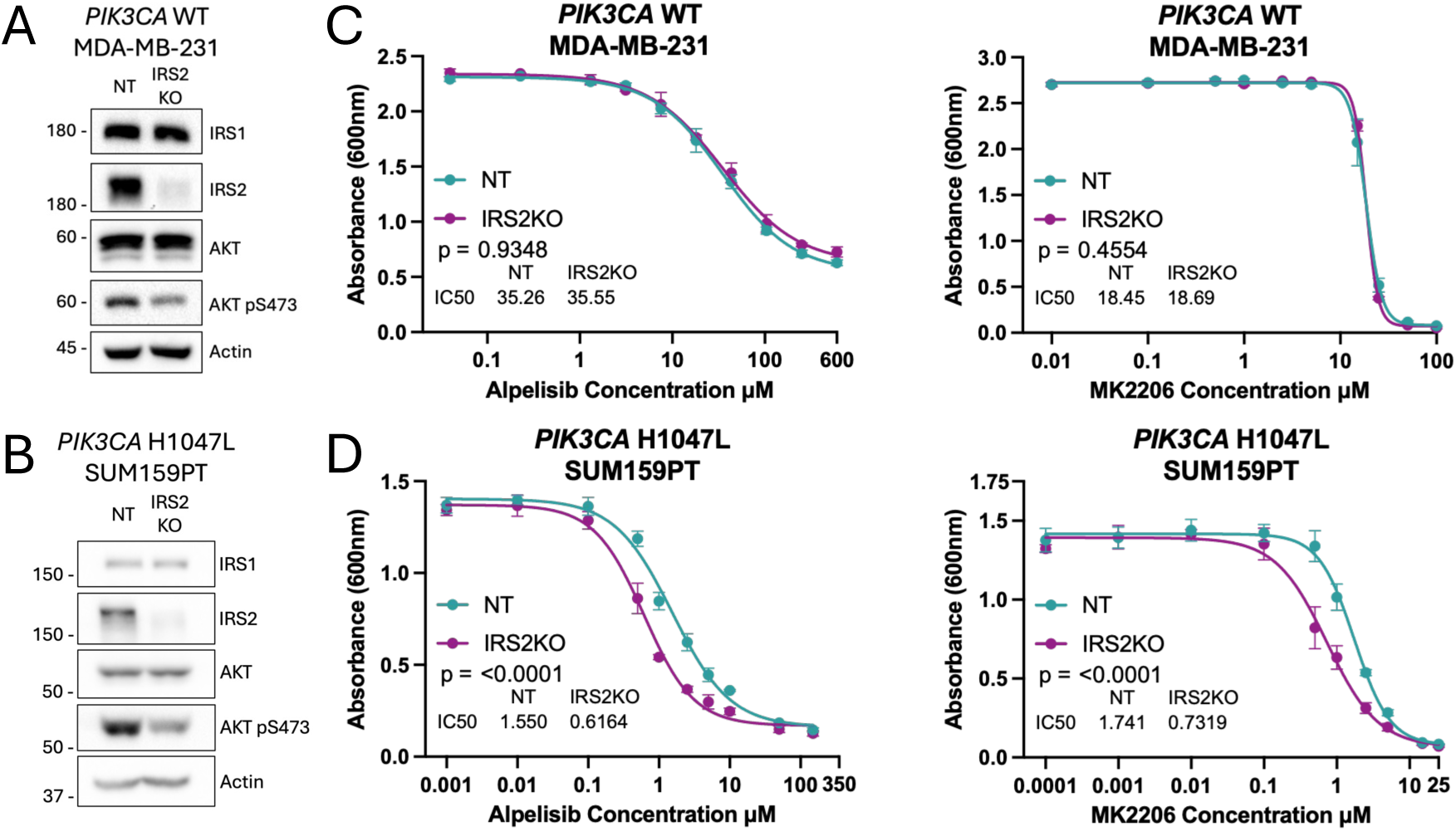
IRS2 promotes resistance to PI3K pathway inhibitors in *PIK3CA* mutant breast cancer cells. (A) MDA-MB-231 cells and (B) SUM159PT cells were treated with non-targeting guide RNA (*NT*) or IRS2-targeting guide RNA (IRS2KO) and cell extracts were analyzed by immunoblot. (C) Viability (live cell population size) of *PIK3CA* WT MDA-MB-231 NT and IRS2KO cells treated with increasing concentrations of alpelisib or MK2206 for 72 hrs, measured by crystal violet staining. (D) Viability of *PIK3CA* mutant SUM159PT NT and IRS2KO cells treated with increasing concentrations of alpelisib or MK2206 for 72 hrs, measured by crystal violet staining. (C and D) Statistical analysis by variable slope (four parameters) nonlinear regression using least squares regression to fit the curve (with asymmetrical confidence intervals calculated for parameters) and the extra sum-of-squares F test to compare IC50 values. Data shown are representative of three independent experiments. See also Figure S4.

To examine further if IRS2-dependent resistance to PI3K pathway inhibition is dependent upon *PIK3CA* mutation alone, experiments were repeated in isogenic MCF10A human mammary epithelial cells with WT *PIK3CA* or heterozygous knock-in of *PIK3CA* E545K or H1047R mutations(4). Control (NT) and IRS2-null (IRS2KO) *PIK3CA* WT, E545K, and H1047R cells were generated using CRISPR-Cas9 gene editing (Fig S5A). For each cell line, two IRS2KO single-cell clones were isolated, maintained separately, and pooled together while plating cells for each viability experiment.

Crystal violet staining cannot determine if differences in cell proliferation or cell death are responsible for variations in cell population size. To determine if the increased sensitivity to PI3K pathway inhibition of *PIK3CA* mutant cells was caused by differences in cell proliferation or cell death, we conducted fluorescence-based and lysis-dependent inference of cell death kinetics (FLICK) assay experiments(55). Briefly, cells were incubated with increasing doses of drug and a fixed concentration of SYTOX Green. SYTOX Green increases in fluorescence upon binding to DNA, is excluded by live cells, and is non-toxic to cells(55). SYTOX green also detects dead cells regardless of the mechanism of cell death, and the signal is proportional to the number of dead cells present(55). Plates are either lysed at the start of the experiment, or are read throughout the drug treatment time course, then lysed and read after 72 hours of drug treatment. Using these readings, it is possible to determine the size of the live and dead cell populations at measured timepoints. These data can be used to calculate relative viability (RV), which is the size of the live cell population relative to the lowest drug dose, and is influenced by both cell proliferation and cell death(56, 57). FLICK can also be used to calculate Lethal Fraction (LF), which is the percent of cells that are dead. Together, these metrics characterize the cytostatic and cytotoxic effects of a drug. A decrease in RV may stem from decreased cell proliferation or increased cell death, or both, which illustrates the importance of measuring LF to determine how a drug is causing a decrease in cell population size.

Consistent with the observations for WT *PIK3CA* expressing MDA-MB-231 cells, there was no difference in relative live cell population size (RV) between control and IRS2-null *PIK3CA* WT MCF10A cells treated with PI3K or AKT inhibitors (Fig 5A,B). In *PIK3CA* WT MCF10A cells, these drugs did induce cell death at high doses, and there was significantly more cell death in IRS2-null cells compared to control cells, as determined through area under curve (AUC) comparison. AUC comparison was utilized because incomplete cell death curves (LF) prevented accurate calculation of IC50 values (see Materials and Methods)(58–60). The observation that differences in cell death (LF) did not cause differences in the relative live cell population sizes (RV) indicates that the differences in cell death between control and IRS2-null cells were modest.

**Figure 5.**
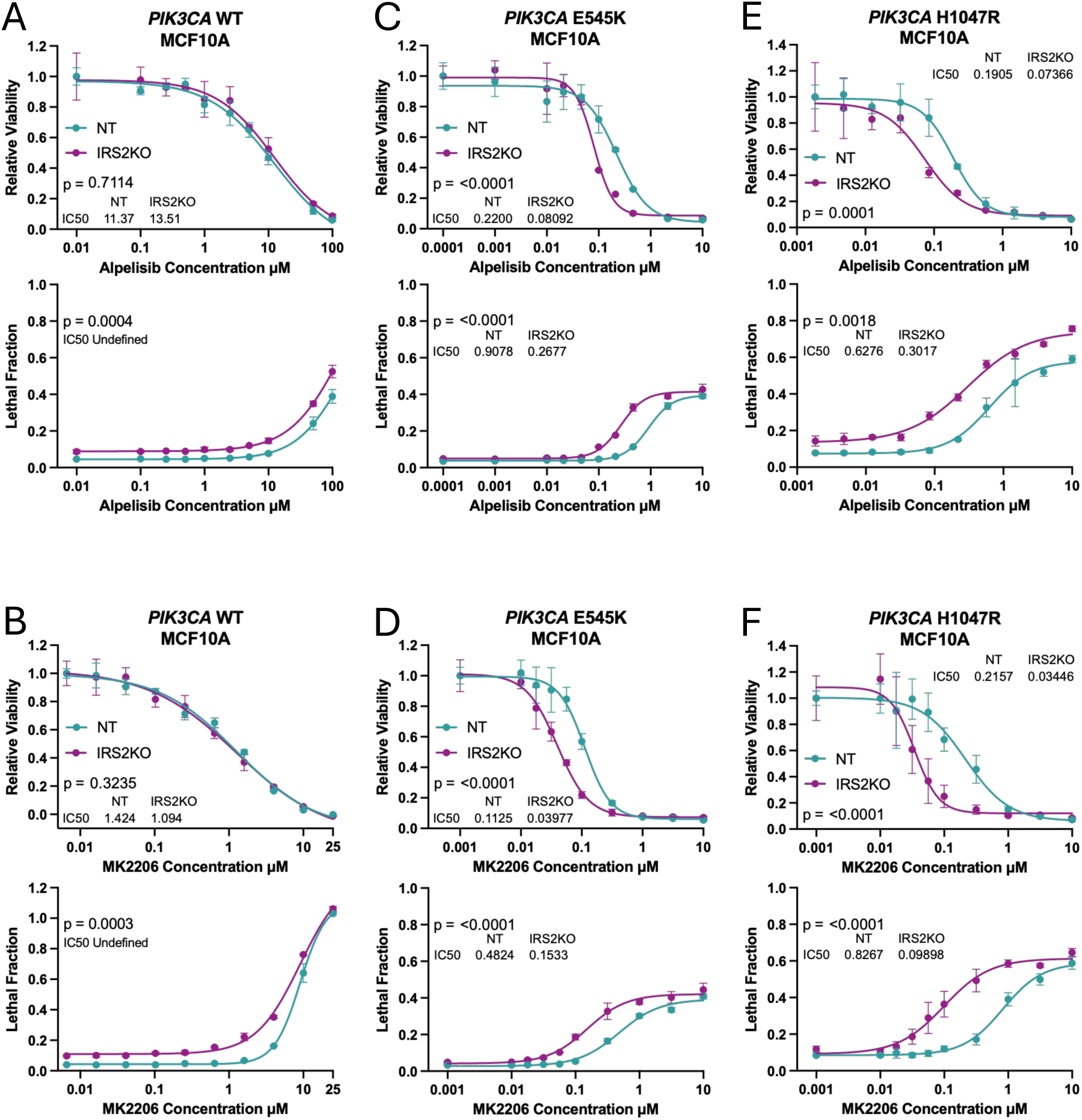
IRS2 depletion reduces cell proliferation and increases cell death in *PIK3CA* mutant mammary epithelial cells in response to PI3K pathway inhibitors. (A,C,E) Relative Viability (top graph) and Lethal Fraction (bottom graph) of NT and IRS2KO *PIK3CA* WT (A), *PIK3CA* E545K (C), and *PIK3CA* H1047R (E) MCF10A cells treated with increasing concentrations of alpelisib for 72 hrs, measured by FLICK assay. (B,D,F) Relative Viability (top) and Lethal Fraction (bottom) of NT and IRS2KO *PIK3CA* WT (B), *PIK3CA* E545K (D), and *PIK3CA* H1047R (F) MCF10A cells treated with increasing concentrations of MK2206 for 72 hrs, measured by FLICK assay. Statistical analysis by variable slope (four parameters) nonlinear regression using least squares regression to fit the curve (with asymmetrical confidence intervals calculated for parameters) and the extra sum-of-squares F test to compare IC50 values for all panels except Lethal Fraction for (A) and (B). Statistical analysis for Lethal Fraction for (A) and (B) by measuring area under curve (AUC) (GraphPad Prism). AUCs were compared using Welch’s t test (two-tailed), utilizing AUC total area and standard error. For AUC comparison of Lethal Fraction for (A) and (B), P values shown are for comparison of data after normalization to lowest drug dose for each subline. P values for original data (shown in graphs) are p<0.0001 (A and B). For all panels, data shown are representative of three independent experiments. See also Figure S5.

In contrast to *PIK3CA* WT cells, both *PIK3CA* E545K and H1047R cells lacking IRS2 were more sensitive to PI3K and AKT inhibition compared to control cells, with IRS2-null cells exhibiting both reduced relative cell population size (RV) and increased cell death (LF) (Fig 5C-F). For IRS2KO cells, RV and LF IC50 values were less than half of control IC50 values for both inhibitors, similar to the increased sensitivity observed in IRS2-null *PIK3CA* mutant SUM159PT (H1047L) cells (Fig 4D). Importantly, in the MCF10A *PIK3CA* mutant cell lines treated with PI3K or AKT inhibitors, differential cell death only partly accounted for differences in relative viability, indicating that ablation of IRS2 also reduced proliferation in these cells.

To determine the impact of IRS2 depletion on cell proliferation and cell death in *PIK3CA* mutant tumor cells treated with PI3K pathway inhibitors, FLICK experiments were conducted using SUM159PT (*PIK3CA* H1047L) IRS2-WT (NT) and IRS2KO breast cancer cells. In these cells, both PI3Kα and AKT inhibition decreased the relative live cell population size (RV) in IRS2KO cells compared to control cells, with IRS2KO IC50 values markedly lower than control IC50 values for both alpelisib and MK2206 (Fig 6A,B). These results are consistent with findings in *PIK3CA* mutant MCF10A cells (Fig 5), and with findings from crystal violet viability experiments in SUM159PT cells (*PIK3CA* H1047L) (Fig 4). Interestingly, while both PI3K and AKT inhibition did induce cell death in the PI3K mutant cells at high doses (increase in LF), there was no difference in the percentage of cells that were alive between control and IRS2KO cells. This result contrasts with the results of MCF10A (*PIK3CA* E545K, H1047R) experiments (Fig 5), in which IRS2-null cells exhibited differences in relative viability (RV) and in cell death (LF). This lack of increased cell death in IRS2-null SUM159PT cells may be due to the presence of additional tumor-promoting mutations in these cells that are absent in MCF10A cells and that compensate for the loss of IRS2-dependent survival signaling. While the mutational background of a given cell line may influence cell survival in the absence of IRS2 expression, experiments in multiple *PIK3CA* mutant cell lines consistently show that IRS2-null cells exhibit increased sensitivity to PI3K pathway inhibition which leads to a reduction in cell proliferation.

**Figure 6.**
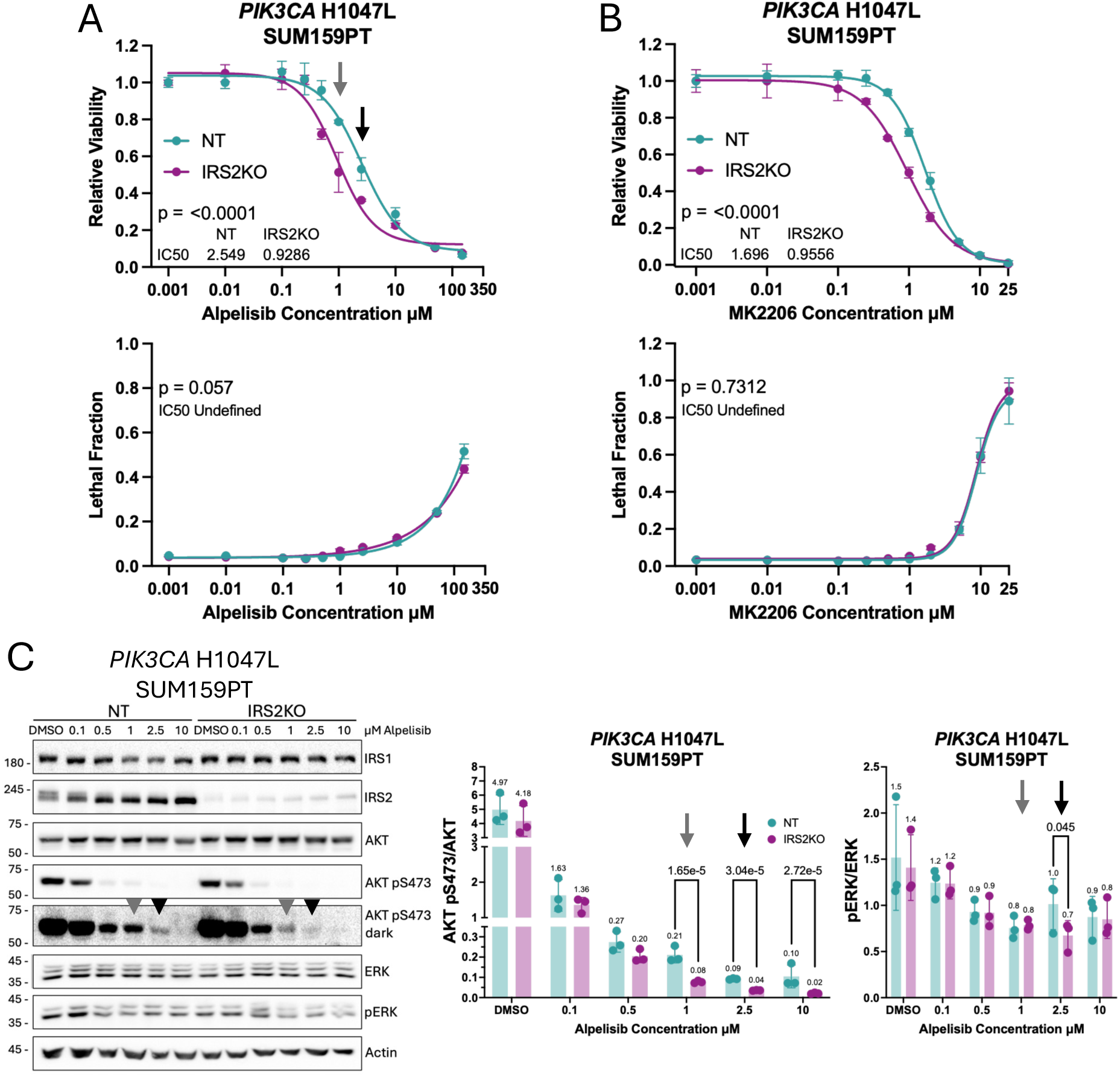
IRS2 sustains PI3K signaling and cell proliferation in *PIK3CA* mutant breast cancer cells treated with PI3K pathway inhibitors. (A and B) Relative Viability (top graph) and Lethal Fraction (bottom graph) of NT and IRS2KO *PIK3CA* H1047L mutant SUM159PT cells treated with increasing concentrations of alpelisib (A) or MK2206 (B) for 72 hrs, measured by FLICK assay. Statistical analysis by variable slope (four parameters) nonlinear regression using least squares regression to fit the curve (with asymmetrical confidence intervals calculated for parameters) and the extra sum-of-squares F test to compare IC50 values for Relative Viability (A and B). Statistical analysis for Lethal Fraction for (A) and (B) by measuring area under curve (AUC) (GraphPad Prism). AUCs were compared using Welch’s t test (two-tailed), utilizing AUC total area and standard error. Data shown are representative of three independent experiments. (C) NT and IRS2KO *PIK3CA* H1047L mutant SUM159PT cells were treated with increasing concentrations of alpelisib for 24 hrs and cell extracts were evaluated by immunoblot. The data shown in the graphs represent the mean +/- SD of three independent experiments. Statistical analysis was performed using a natural-log transformation followed by ordinary least squares (OLS) ANOVA on log(ratio), with fixed effects for genotype (NT vs IRS2KO), dose (categorical: 0 (DMSO), 0.1, 0.5, 1, 2.5, 10), their interaction, and a gel blocking factor, according to the model: log(ratio) ∼ genotype × dose + gel. Pre-specified comparisons between NT and IRS2KO were conducted within each dose on the log scale. Because these comparisons were planned a priori, no multiplicity adjustments were applied (P values shown).

### IRS2 promotes resistance to PI3K inhibition by activating PI3K signaling

To explore the potential mechanism(s) for IRS2-dependent resistance to PI3K pathway inhibition, we compared cell viability and IRS2-dependent signaling pathway activity during PI3Kα inhibition (Fig 6C). PI3K/AKT and ERK signaling were assessed because IRS2 is known to activate both pathways and both pathways can regulate cell proliferation(61). Compared to control cells, there was a significant reduction in PI3K/AKT signaling (AKT pS473) in IRS2-null cells at doses of alpelisib that elicited divergent cell viability (1μM, 2.5μM) (Fig 6A,C). There was also reduced ERK signaling in IRS2-null cells as measured by ERK phosphorylation at 2.5μM, although the difference between IRS2KO and control cells was smaller than for PI3K signaling.

To further probe the mechanism of IRS2-mediated resistance to PI3K pathway inhibition, IRS2-null SUM159PT cells (*PIK3CA* H1047L) were transduced to express empty vector (EV) (control), wild-type IRS2 (IRS2-WT), or IRS2-Y5F, which is a full-length mutant form of IRS2 which includes five tyrosine to phenylalanine mutations that block IRS2 recruitment of the p85 subunit of PI3K (Fig 7A)(62). IRS2 Y5F is useful because it only interrupts the ability of IRS2 to activate PI3K, it does not affect ERK signaling(62). Both IRS2 WT and IRS2 Y5F were expressed at equivalent levels (Fig 7A, Fig S6A). *PIK3CA* mutant cells rescued with IRS2-WT expression were more resistant to PI3Kα inhibition compared to EV control cells (Fig 7B). The magnitude of this effect was consistent with prior experiments in SUM159PT NT and IRS2KO cells (Fig 4D, Fig 6A). Also consistent with these prior experiments, the restoration of IRS2-WT expression did not impact cell death. Cells expressing IRS2-Y5F retained a greater sensitivity to PI3Kα inhibition, similar to EV control cells. This finding supports that the ability of IRS2 to recruit and signal through PI3K is essential for IRS2-dependent resistance to PI3K pathway inhibition.

**Figure 7.**
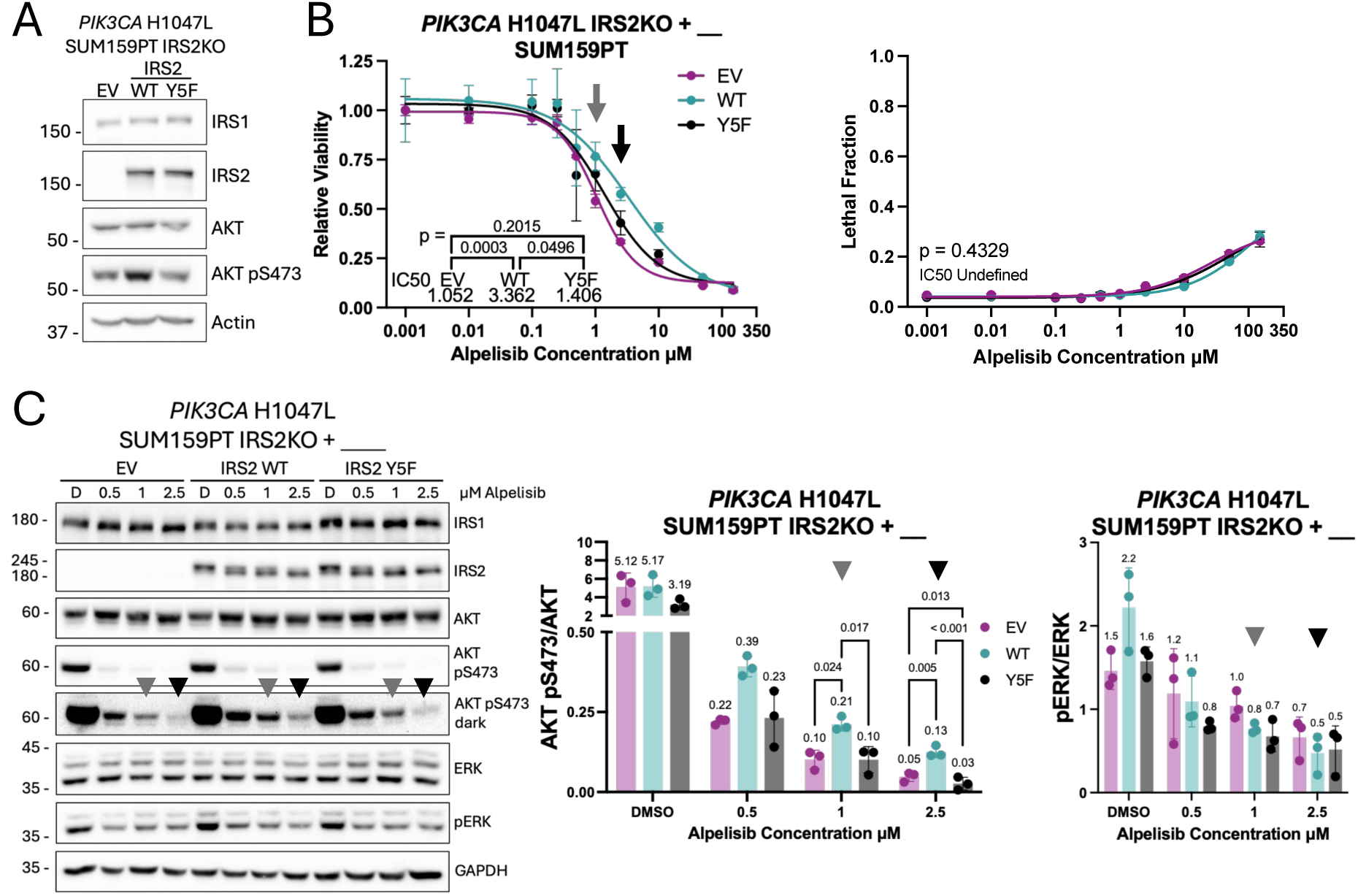
IRS2-dependent PI3K activation confers resistance to PI3K inhibition in *PIK3CA* mutant breast cancer. (A) Cell extracts from *PIK3CA* H1047L mutant SUM159PT IRS2KO cells expressing empty vector (EV), IRS2-WT or IRS2-Y5F were analyzed by immunoblot. (B) Relative Viability (left graph) and Lethal Fraction (right graph) of *PIK3CA* H1047L mutant SUM159PT IRS2KO cells expressing empty vector (EV), IRS2-WT or IRS2-Y5F treated with increasing concentrations of alpelisib for 72 hrs, measured by FLICK assay. Statistical analysis of Relative Viability by variable slope (four parameters) nonlinear regression using least squares regression to fit the curve (with asymmetrical confidence intervals calculated for parameters) and the extra sum-of-squares F test to compare pairs of IC50 values. Statistical analysis for Lethal Fraction by measuring area under curve (AUC) (GraphPad Prism). AUCs were compared using One-Way ANOVA followed by the Tukey test for multiple comparisons to compare all treatments, utilizing AUC total area and standard error. Data shown are representative of three independent experiments. (C) *PIK3CA* H1047L mutant SUM159PT IRS2KO cells expressing empty vector (EV), IRS2-WT or IRS2-Y5F were treated with increasing concentrations of alpelisib for 24 hrs and cell extracts were analyzed by immunoblot (D = DMSO). The data shown in the graphs represent the mean +/- SD of three independent experiments. Statistical analysis was performed using a natural-log transformation followed by ordinary least squares (OLS) ANOVA on log(ratio), with fixed effects for genotype (EV vs. IRS2-WT vs. IRS2-Y5F), dose (categorical: 0 (DMSO), 0.5, 1, 2.5), their interaction, and a gel blocking factor, according to the model: log(ratio) ∼ genotype × dose + gel. Pre-specified comparisons between each genotype pair within each dose were conducted on the log scale. Because these comparisons were planned a priori, no multiplicity adjustments were applied (P values shown). See also Figure S6.

PI3Kα inhibition significantly reduced PI3K signaling in EV control and IRS2-Y5F expressing cells compared to IRS2-WT cells at doses which also yielded differences in viability (1μM, 2.5μM) (Fig 7B,C). In contrast, there were no differences in ERK activity between EV control, IRS2-WT, and IRS2-Y5F expressing cells (Fig 7C). Collectively, the inability of IRS2-Y5F to restore resistance to PI3K inhibition, coupled with reductions in PI3K signaling which correspond with increased sensitivity to alpelisib, indicate that IRS2-dependent PI3K signaling mediates resistance to PI3K inhibition in *PIK3CA* mutant cells.

### IRS2 expression influences PI3K pathway inhibitor sensitivity across *PIK3CA* mutant cancers

To determine if IRS2 confers resistance to PI3K pathway inhibitors in a wider cancer context, the Cancer Dependency Map (DepMap) was used to generate correlations between IRS1 or IRS2 mRNA or protein abundance and PI3Kα inhibitor sensitivity (alpelisib) for all *PIK3CA* mutant cell lines arising from solid tumors (Fig 8)(43). No relationship between IRS1 mRNA or protein levels and PI3Kα inhibitor sensitivity was observed (Fig 8A,B). However, *IRS2* mRNA abundance correlated with PI3Kα inhibitor sensitivity, with higher levels of *IRS2* associated with increased resistance to PI3Kα inhibition (as measured by increased AUC) (Fig 8A). This correlation was also present for IRS2 protein levels (Fig 8B). To interrogate if this association was unique to *PIK3CA* mutant cell lines, *IRS1* and *IRS2* mRNA was correlated to PI3Kα inhibitor sensitivity in *PIK3CA* wild-type cells arising from all solid tumors (Fig S7A). *IRS1* did not correlate with drug sensitivity, whereas *IRS2* showed a modest positive and significant association with resistance to PI3Kα inhibition. However, linear regression on data from *PIK3CA* mutant cells exhibited a steeper slope and higher Pearson correlation coefficient compared to the *IRS2* - alpelisib sensitivity plot from wild-type cells, indicating that *IRS2* mRNA abundance had a larger impact on alpelisib sensitivity in *PIK3CA* mutant cells than in wild-type cells. A similar trend was observed for cell line sensitivity to the AKT inhibitor MK2206, with only *IRS2* being associated with drug sensitivity in *PIK3CA* mutant cells (Fig S7B).

**Figure 8.**
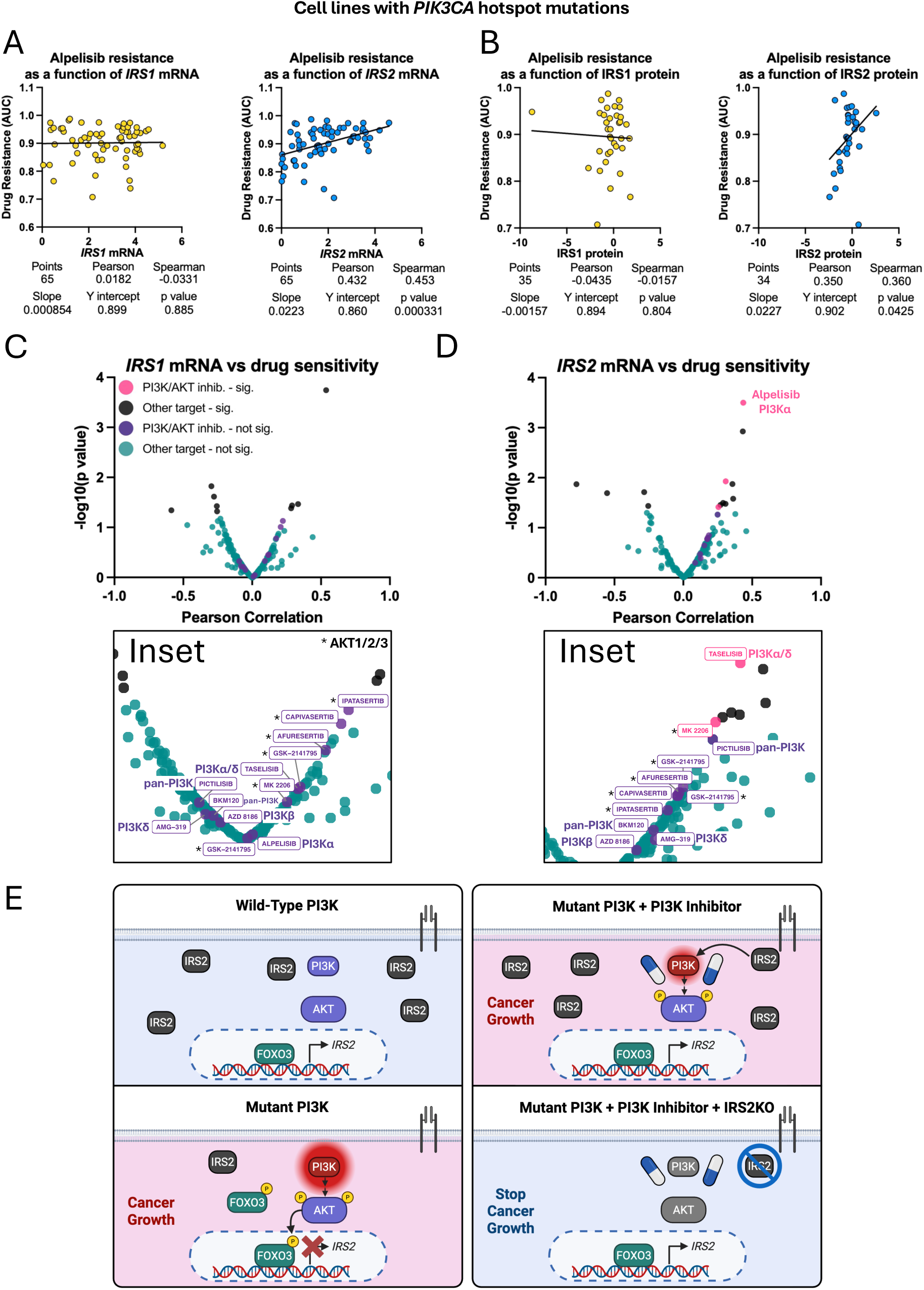
IRS2 expression positively correlates with resistance to PI3K pathway inhibitors in *PIK3CA* mutant cancer cell lines. (A and B) Correlation between IRS1 and IRS2 mRNA (A) or protein (B) and sensitivity to alpelisib in cancer cell lines with *PIK3CA* hotspot mutations (excluding myeloid and lymphoid-derived cell lines) in the DepMap dependency dataset. P values shown are from simple linear regression, p<0.05 indicates that the slope is significantly different from zero. mRNA abundance shown is log2(TPM+1). (C and D) Correlation between *IRS1* (C) and *IRS2* (D) mRNA (DepMap) and sensitivity to drugs tested in the Sanger Genomics of Drug Sensitivity in Cancer 2 (GDSC2) drug panel in *PIK3CA* mutant cancer cell lines (excluding myeloid and lymphoid-derived cell lines) (accessed using the DepMap dependency dataset). Drug sensitivity was determined by AUC. GSK-2141795 was tested twice in GDSC2 and both results are plotted. Pearson correlation of mRNA abundance (log2(TPM+1)) and drug sensitivity (AUC). P values (y axis) are from simple linear regression of mRNA abundance vs. drug sensitivity. Significance defined as p<0.05. Abbreviations, Sig. for significant, inhib. for inhibitor. Drug names and targets listed (inset). Pan-PI3K – inhibitor of PI3Kα/β/δ/𝛾. (E) Model for IRS2 conferring resistance to PI3K pathway inhibition in PI3K-mutant breast cancer. *PIK3CA* mutations activate PI3K signaling to inactivate FOXO3, leading to reduced IRS2 mRNA and protein abundance. PI3K inhibition leads to FOXO3-driven increases in IRS2. During PI3K inhibition, IRS2 sustains PI3K signaling and promotes *PIK3CA* mutant cancer cell viability. Ablation of IRS2 during PI3K inhibition lowers PI3K signaling, leading to reduced cancer cell population growth (Created with BioRender.com). See also Figure S7.

We expanded these analyses by assessing the correlation of *IRS1* and *IRS2* expression with sensitivity to a broad panel of cancer targeting drugs, that included additional agents targeting the PI3K pathway, in *PIK3CA* mutant cancer cell lines arising from solid tumors using DepMap (Fig 8C,D)(43, 63, 64). Higher *IRS2* expression significantly correlated with resistance to alpelisib (a PI3Kα-selective inhibitor) and MK2206 (an AKT1/2/3 inhibitor), which were both used in our studies (Fig 8D). A significant association with resistance was also observed for taselisib, a PI3Kα/δ-selective inhibitor. The only PI3Kα-selective inhibitors included in the drug panel, alpelisib and taselisib, had the first and third highest correlation with *IRS2* expression of all compounds tested, respectively. While it is important to test this finding with additional PI3Kα-selective inhibitors, this raises the possibility that IRS2 depletion would be most sensitizing to PI3Kα inhibitors, several of which have been approved clinically for the treatment of *PIK3CA* mutant breast cancer(16, 17). In contrast, *IRS1* expression was not significantly associated with resistance to any of the PI3K or AKT inhibitors tested (Fig 8C). Together, these observations support that IRS2 promotes resistance to PI3K pathway inhibitors in *PIK3CA* mutant cancers from a variety of tissues (Fig 8E).

## Discussion

In this study, we establish a compensatory feedback mechanism by which PI3K pathway inhibitors increase IRS2 abundance and demonstrate the role of IRS2 in promoting resistance to these drugs. IRS2 mRNA and protein levels are lower in *PIK3CA* mutant breast cancer patient tumors and cell culture models of breast cancer. This phenotype involves PI3K-dependent inactivation of the transcription factor FOXO3 and is reversed by PI3K or AKT inhibition. In contrast, IRS1 expression is not altered by *PIK3CA* mutations. IRS2 expression confers resistance to PI3K pathway inhibitors in *PIK3CA* mutant, but not WT breast cancer, by sustaining PI3K signaling. Increased IRS2 abundance correlates significantly with resistance to many PI3K pathway inhibitors across cancer cell lines arising from diverse tissues. Our findings have clinical relevance given the frequency of *PIK3CA* mutations in cancer, the importance of PI3K signaling reactivation in resistance to PI3K pathway inhibitors, and the identification of a new target to address the challenges associated with prior efforts to block PI3K pathway reactivation during PI3K inhibition.

Analysis of publicly available data revealed that IRS2 mRNA and protein abundance is decreased in *PIK3CA* mutant human breast tumors and breast cancer cell lines. This reduced level of IRS2 expression may seem counterintuitive given the previously demonstrated role of IRS2 in promoting tumor progression and metastasis and the more aggressive nature of *PIK3CA* mutant tumors. However, although IRS2 is expressed at lower levels in these cells, it remains functionally important. For example, in SUM159PT cells (*PIK3CA* H1047L), stimulation with insulin or IGF-1 further enhances PI3K pathway activity, and IRS2 is required for efficient invasion and breast cancer stem cell self-renewal(37, 38, 50). *PIK3CA* mutant cells may only require lower levels of IRS2 because the majority of the necessary PI3K signaling is provided through mutant PI3K. However, when *PIK3CA* mutant cells experience reduced PI3K signaling due to PI3K inhibition, compensatory pathways for PI3K activation are necessary and elevated levels of IRS2 are required to maintain adequate PI3K signaling for continued viability. The insulin-PI3K-FOXO regulatory feedback pathway that plays an important role in controlling normal metabolic homeostasis is highjacked in *PIK3CA* mutant tumor cells to sustain viability in response to targeted pathway inhibitors. One aspect of note from our study is that IRS2 ablation sensitizes *PIK3CA* mutant cells, but not *PIK3CA* wild-type cells, to PI3K pathway inhibitors. In line with this finding, our results, as well as prior literature, indicate that *PIK3CA* mutant cancer cells are more sensitive to PI3K pathway inhibitors compared to *PIK3CA* wild-type cancer cells(14, 65–67). This may be due to an increased dependence on the upregulation of compensatory PI3K signaling mechanisms, such as IRS2, during PI3K pathway inhibition in *PIK3CA* mutant cancer cells compared to *PIK3CA* WT cells. This also explains why a previous effort to identify mechanisms of resistance to PI3K inhibition that did not evaluate *PIK3CA* wild-type and mutant cells separately failed to identify IRS2 as a driver of PI3K inhibitor resistance, as IRS2 ablation sensitizes only *PIK3CA* mutant cells to PI3K pathway inhibition(68–70).

Our findings on the role of IRS2 in promoting PI3K pathway inhibitor resistance suggest that other molecules associated with IRS2 function and signaling could also be involved. For example, several groups have reported that PI3K pathway inhibitors cause IGF1R upregulation, which leads to re-activation of PI3K signaling and drug resistance(29, 30, 32). Other studies implicate MYC in resistance to PI3K pathway inhibitors(26, 71). Previous work from our lab reported that IGF1R signals through IRS2, but not IRS1, to stabilize MYC and support breast cancer stem cell activity, suggesting that this signaling axis could also be responsible for the differential contributions of IRS1 and IRS2 to PI3K pathway inhibitor resistance that we observed(38). IRS2-dependent MYC stabilization could also explain why IRS2 ablation coupled with PI3K pathway inhibition limits cell proliferation. Given the role of IRS2 in activating PI3K and AKT, which drive cell survival, and the previously described role of IRS2 in promoting cell survival, it was somewhat surprising that IRS2 loss coupled with PI3K pathway inhibition increased cell death in MCF10A *PIK3CA* mutant cells, but not in *PIK3CA* mutant SUM159PT cells(72). However, these knock-in MCF10A cells carry no mutations other than mutant *PIK3CA*, whereas SUM159PT cells harbor mutations in *PIK3CA* as well as *TP53* and *HRAS*, which may promote cell survival and be responsible for this divergent phenotype(73). Future analysis of a broader number of *PIK3CA* mutant tumor cell lines will be necessary to determine the genetic background that compensates for the loss of IRS2-dependent cell survival signaling during PI3K inhibition.

From a clinical perspective, the frequency and severity of adverse events caused by PI3K pathway inhibitors limits their utility. In a phase 3 clinical trial of alpelisib, the PI3Kα specific inhibitor used in our study, over 60% of patients had to take a reduced dose due to side effects, and 25% of patients had to stop the treatment altogether(14). Other therapies targeting this pathway also suffer from frequent dose-limiting side effects(15, 74). Treatment-induced hyperglycemia and gastrointestinal symptoms, which constitute some of the most frequent severe side effects, stem from inhibition of PI3K in non-tumor tissues(75). One approach to reducing side effects during PI3K inhibition is for patients to limit hyperglycemia by adhering to a low carbohydrate, ketogenic diet. While this approach attenuates PI3K-inhibitor induced hyperglycemia and hyperinsulinemia in mouse models of cancer and is being tested clinically, it is not a perfect solution, as patients often have difficulty adhering to a strict ketogenic diet(21, 76, 77). Use of an SGLT2 inhibitor to decrease hyperglycemia during PI3K inhibition is also being evaluated(77). While carbohydrate restriction and SGLT2 inhibition may minimize hyperglycemia in patients, these approaches are unlikely to reduce gastrointestinal side effects because these side effects are due to PI3K inhibition in these tissues, not hyperglycemia.

Sensitizing *PIK3CA* mutant cancers to PI3K pathway inhibitors through depletion of IRS2 could be a more promising approach to improve clinical outcomes because patients would require lower drug doses (Fig 8E). This would limit all on-target and off-target side effects, rather than only those due to hyperglycemia, while allowing IRS1 to signal normally for metabolic regulation. This metabolic redundancy may lead to a better safety profile for IRS2 depletion coupled with PI3K inhibition compared to blocking PI3K pathway reactivation during PI3K inhibition by targeting IGF1R or mTOR, as side effects from these combinations limit drug tolerability(33, 78–80). IRS2 has received limited attention as a potential therapeutic target because it is a cytoplasmic adaptor protein with no active site and is therefore not amenable to inhibition with antibodies or small molecules, respectively. While one IRS-targeting agent has been reported, it has pleiotropic effects that suppress both IRS1 and IRS2 indirectly and has not been pursued clinically(81, 82). We recently demonstrated that small interfering RNAs (siRNAs) conjugated to an albumin binding dendrimer selectively suppress *Irs2*, but not *Irs1*, expression in mammary tumors without causing hyperglycemia(83). With the emergence of these RNA-based gene silencing drug modalities, depletion of IRS2 in combination with a PI3K/AKT inhibitor may be a feasible future approach for the treatment of PI3K mutant cancer.

### Experimental Procedures

#### Cells and Cell Culture

MCF10A cells (*PIK3CA* wild-type and *PIK3CA* knock-in mutant cells) were a gift from Dr. Ben Ho Park (Vanderbilt). *PIK3CA* mutant MCF10A cells were maintained in DMEM/F12 (Gibco) containing 5% horse serum (Gibco), 0.5 μg/mL hydrocortisone (Sigma-Aldrich), 100ng/mL cholera toxin (Sigma-Aldrich), 10 μg/mL insulin (Sigma-Aldrich), and 100U/mL Penicillin-Streptomycin (Gibco). *PIK3CA* wild-type MCF10A cells were maintained in identical media that also contained 20ng/mL human EGF (Sigma-Aldrich). *PIK3CA* wild-type and mutant MCF10A cells were plated in the same media for 18-24 hours prior to starting experiments. SUM159PT cells were obtained from BioIVT and were grown in F12 media (Gibco) containing 5% FBS (Cytiva), 25mM HEPES (Sigma-Aldrich), 5 µg/mL insulin (Sigma-Aldrich) and 1 µg/mL hydrocortisone (Sigma-Aldrich). MDA-MB-231, T47D, and MCF7 human breast carcinoma cells were obtained from the ATCC Cell Biology Collection. MDA-MB-231 and T47D cells were grown in RPMI (Gibco) containing 10% FBS (Cytiva). MCF7 cells were grown in MEM (Gibco) containing 10% FBS (Cytiva) and 11.2μg/mL insulin (Sigma-Aldrich). UACC812 and HCC1806 human breast carcinoma cells were a gift from Dr. Dohoon Kim (UMass Chan). UACC812 cells were maintained in DMEM 4.5g/L glucose (Gibco) containing 100U/mL Penicillin-Streptomycin (Gibco), 2mM L-Glutamine (Gibco), and 10% FBS (Cytiva). HCC1806 cells were grown in RPMI (Gibco) containing 10% FBS (Cytiva). BT20 human breast carcinoma cells were a gift from Dr. Michael Lee (UMass Chan) and were grown in MEM (Gibco) containing 10% FBS (Cytiva). Cells were stimulated with human insulin-like growth factor 1 (IGF1) as described in figure legends (Sigma-Aldrich). All cells were screened regularly and tested negative for mycoplasma by PCR (Abm).

IRS2 (*IRS2^-/-^*) knock-out and control (NT) cells were generated by CRISPR/Cas9-mediated gene editing by electroporation of a ribonucleoprotein (RNP) complex of CRISPR-gRNA and Cas9 protein prepared according to manufacturer’s protocol (IDT). Cas9 used was Alt-R™ S.p. Cas9 Nuclease V3 (IDT). Guide RNA used: sgIRS2, CAA GGA CGA GTA CTT CGC CG (IDT Hs.Cas9.IRS2.1.AA). The control non-targeting guide RNA (sgNT) was purchased from IDT (Alt-R™ Cas9 Negative Control crRNA #1). Electroporation of RNP complex was done using nucleofector kit V (Lonza) and Nucleofector II electroporation machine (Amaxa, Program X-013 (SUM159PT and MDA-MB-231), Program T-020 (MCF10A)).

FOXO3 knock-down and control (shGFP) cells were generated using simple hairpin shRNAs in the pLKO.1 lentiviral vector designed by The RNAi Consortium (TRC) (UMass Chan RNAi Core Facility). shRNA used: shFOXO3 #1 - Clone Id: TRCN0000040098, mature antisense TTG TGT CAG TTT GAG GGT CTG; shFOXO3 #2 - Clone Id: TRCN0000040102, mature antisense ATG AAT CGA CTA TGC AGT GAC; shFOXO3 #3 - Clone Id: TRCN0000010334, mature antisense GGT ATA CTT GTT GCT ATT GT; shGFP control - Catalog ID: RHS4459. shRNAs were provided by the UMass Chan RNAi Core Facility as viral supernatants. MCF10A (*PIK3CA* H1047R) cells were infected for 24h with virus in the presence of 8 μg/mL polybrene (Sigma-Aldrich) and selected with 1 µg/mL puromycin (Gold Biotechnology).

#### Plasmids

Human IRS2 was a kind gift from Adrian Lee (University of Pittsburgh, Pittsburgh, PA). cDNA was subcloned into the pCDH-CMV-MCS-EF1-puro lentiviral vector (System Bioscience) with a C-terminal FLAG tag(84). The IRS2-Y5F mutant construct was generated by Genewiz and subcloned into the pCDH-CMV-MCS-EF1-puro lentiviral vector (System Bioscience) with a C-terminal FLAG tag(62). Empty vector (EV) control plasmid was the pCDH-CMV-MCS-EF1-puro lentiviral vector (System Bioscience). Lentiviruses were generated by co-transfection of IRS2 plasmids and packaging plasmids (pMD2.G and psPAX2) into HEK293FT cells using lipofectamine 3000 according to the manufacturer’s instruction (Invitrogen). Cells were infected for 24 h with virus in the presence of 8 μg/mL polybrene (Sigma-Aldrich) and selected with 2 µg/mL puromycin (Gold Biotechnology).

#### Drugs

Alpelisib was prepared at 150mM in DMSO prior to diluting for experiments (MedChemExpress). MK2206 was prepared at 25mM in DMSO prior to diluting for experiments (MedChemExpress).

#### Crystal Violet Viability Assay

Cells were split on day 1, then trypsinized, counted, and plated in 96 well plates on day 2. Cells were not plated in perimeter wells; these were filled with PBS. On day 3, media was removed, and fresh media containing drugs was added. Plates were incubated for 72 hours, then media was removed and cells were fixed in paraformaldehyde (4% in PBS) for 40 minutes (Boston Bioproducts). Paraformaldehyde was removed and cells were stained with crystal violet (0.2%) for 70 minutes (Sigma-Aldrich). Following staining, wells were washed three times with water and plates were allowed to dry. Once plates were dry, 1% SDS was added to the wells and plates were agitated for 30 minutes to dissolve crystal violet. Absorbance was measured using a microplate reader (Promega GloMax Explorer, software v.4.0.0., absorbance 600nm).

#### FLICK Viability Assay

The FLICK assay was performed as described previously(55). Briefly, cells were incubated with increasing doses of drug and a fixed concentration of SYTOX Green (Invitrogen). Plates were either lysed at the start of the experiment, or were read throughout the drug treatment time course, then lysed and read after 72 hours of drug treatment. Plates were lysed with 0.1% Triton X-100 (final concentration) (BioRad). Black-walled, optical bottom, 96-well microplates were used for these experiments (Corning, Greiner Bio-One, Thermo Scientific). Plates were read with a microplate reader (Ex: 485nm/20nm, Em: 528nm/20nm) (BioTek Synergy HT microplate reader, software v.3.11.19. Optics: Bottom, Gain: 55, Read speed: normal). This microplate reader was suitable for the FLICK assay because the gain can be set manually, which is crucial for accuracy in repeated measurements of the same plate throughout a time course.

#### Immunoblotting

For whole-cell extracts, cells were solubilized at 4 °C in lysis buffer (20 mM Tris buffer, pH 7.4, 1% Nonidet P-40, 0.137 M NaCl, 10% glycerol) containing phosphatase inhibitors (Roche) and protease inhibitors (Roche). Cell extracts containing equivalent amounts of protein were resolved by SDS-PAGE and transferred to nitrocellulose membranes. Membranes were blocked using 5% non-fat milk in 1x TBST (50 mM Tris, pH 7.5, 150 mM NaCl, 0.01% Tween 20). Primary antibody incubations were carried out in 5% bovine serum albumin (BSA) or 5% non-fat milk in 1X TBST overnight at 4 °C. Primary antibodies used in this study include: IRS1 (Cell Signaling Technology [CST] #95816; Bethyl Laboratories #A301-158A, sold by ThermoFisher Scientific), IRS2 (CST #4502), GAPDH (Santa Cruz Biotechnology #sc-32233), phospho-AKT S473 (CST #9271), AKT (CST #9272), α-tubulin (CST #3873), phospho-ERK (CST #9101), ERK (CST #9102), Actin (Invitrogen # MA5-15739), FOXO1 (CST #2880), FOXO3a (CST #12829), phospho-FOXO1 S256 (CST #9461), phospho-FOXO3a S253 (CST #13129), and Lamin A/C (Santa Cruz Biotechnology #sc-376248). Secondary antibody (Jackson ImmunoResearch Peroxidase AffiniPure Goat Anti-Rabbit IgG (H+L) #111-035-144 and Goat Anti-Mouse IgG (H+L) #115-035-146) incubations were carried out in 5% non-fat milk in 1X TBST for 1 h at room temperature. Bands were detected by chemiluminescence using a ChemiDoc XRS+ system (Bio-Rad), using Clarity Western ECL Substrate (BioRad) or SuperSignal™ West Femto Maximum Sensitivity Substrate (Thermo Scientific).

For immunoblot quantification, the signals from the protein of interest were normalized with that of the loading control, and the signals from phospho-specific antibodies were normalized to the total protein of interest using Image J (v.1.52) or BioRad Image Lab (v.6.1.0 build 7). Only bands that produced a signal which did not saturate the ChemiDox XRS+ detector were used for quantification. All immunoblotting images used for quantification were 16-bit TIFF files.

#### Nuclear / Cytoplasmic Fractionation

Nuclear and cytoplasmic extraction was carried out using the NE-PER™ Nuclear and Cytoplasmic Extraction Reagents (Thermo Scientific) according to manufacturer’s protocol, with one modification. Following cytoplasmic extraction, (nuclear) pellets were washed once with cold 1X PBS, briefly resuspended, then centrifuged at maximum speed in a microcentrifuge for 1 minute. PBS was then completely removed prior to resuspending the pellets in ice-cold nuclear extraction reagent (NER). Extracts were then analyzed by immunoblotting as described.

#### Quantitative Polymerase Chain Reaction (qPCR)

RNA was extracted using the RNA-easy kit (Qiagen). cDNA was synthesized using an AzuraQuant cDNA synthesis kit (Azura Genomics). qPCR was performed in an Applied Biosystems QuantStudio 6 Flex apparatus using AzuraView GreenFast qPCR Blue Mix (Azura Genomics). The delta-delta Ct method was used to determine relative mRNA expression (r18s used for normalization)(85). Primers used were as follows: h*IRS1*-Fwd 5’-CAA CTG GAC ATC ACA GCA GAA-3’; h*IRS1*-Rev 5’-ACT GAA ATG GAT GCA TCG TAC C-3’; h*IRS2*-Fwd 5’-CCA CCA TCG TGA AAG AGT GA-3’; h*IRS2*-Rev 5’-CAG AGT CCA CAG ATG TTT CCA A-3’; r18s-Fwd 5’-GTC GCT CGC TCC TCT CCT ACT-3’; r18s-Rev 5’-TCT GAT AAA TGC ACG CAT CCC-3’. The average delta Ct for the control condition (WT or WT treated with DMSO) across all replicates was used to calculate delta-delta Ct values. This enables calculation of relative mRNA expression values for all conditions and replicates while also retaining and displaying the variability in the control condition for all replicates. 2^-ΔΔCT^ (relative expression) was then calculated for all conditions and replicates. All relative expression values were then normalized to the average of the three control condition relative expression values so that the mean of the controls was equal to 1.

### Bioinformatics and Statistical Analysis

#### Clinical Data in Violin Plots

Clinical data shown in Figure 1A,B, Figure S1A-E, and Figure S3A were derived from the Molecular Taxonomy of Breast Cancer International Consortium (METABRIC) dataset, accessed using the cBioPortalData (version 2.16.0) R package(39, 86). Gene expression measurements (log2 intensity values) derived from the Illumina HT-12 v3 microarray platform were retrieved for 1,980 total breast cancer samples. Samples were grouped according to *PIK3CA* genomic status into mutant (n = 795) and wild-type (n = 1,185) cohorts. Breast cancer subtypes were defined as Luminal A&B (ER+ and HER2-), Triple-negative (ER-, PR-, HER2-), and HER2 enriched (HER2+) (ER = estrogen receptor. HER2 = human epidermal growth factor receptor 2)(87). Data shown in Figure 1C,D were derived from the Clinical Proteomic Tumor Analysis Consortium (CPTAC) dataset, accessed via https://kb.linkedomics.org/(42). For each gene, expression distributions were assessed for normality using the Shapiro–Wilk test(88). Group-wise expression patterns were visualized using combined violin and box plots generated with ggplot2(version 3.5.2) and ggpubr(version 0.6.1)(89, 90) Statistical comparisons between *PIK3CA* mutant and wild-type groups were performed using the Wilcoxon rank-sum test(91). All statistical analyses and figure generation were performed in R (version 4.4.1)(92).

#### DepMap-Derived Data

Data shown in Figure 2G were from Batch corrected Expression Public 24Q2 dataset (DepMap: The Cancer Dependency Map Project at Broad Institute, depmap.org)(43). Data shown in Figure 2H were from RPPA500 MCLP (DepMap). For Figure 8, data shown is from all cell lines, excluding myeloid and lymphoid-derived cell lines. All mRNA data (Figure 8A,C,D, Figure S7A,B) is from Expression Public 25Q3 (IRS1 or IRS2 log2(TPM+1)). Protein expression for Figure 8B is from Relative Protein Expression Harmonized MS CCLE Gygi (DepMap). Drug sensitivity association results for all panels of Figure 8 and Figure S7 are from Sanger Genomics of Drug Sensitivity in Cancer 2 (GDSC2) (1053 - MK2206, 1560 - alpelisib)(accessed using DepMap)(63, 64). In Figure 8C,D, only drugs that specifically inhibited PI3K or AKT (and did not inhibit other kinases) were labeled as PI3K/AKT inhibitors and were colored pink or purple. PI3K and AKT isoform-specific inhibition for drugs labeled PI3K/AKT inhibitors was determined according to prior literature(93, 94). All volcano plot analyses and figures were generated in R (version 4.4.1) using ggplot2(version 3.5.2) for plotting and ggrepel(version 0.9.6) for non-overlapping labels(89, 92, 95).

#### Statistical Analysis of Cell Viability

IC50 values reported throughout the paper are relative IC50 values. IC50 values and p values from comparison of IC50 values are only provided when sufficient data are present to accurately estimate IC50 values. Mathematical modeling shows that a cell viability curve requires at least two drug concentrations below and two drug concentrations above the linear portion of the curve (the straight line that includes the IC50) to accurately calculate the IC50 and therefore to accurately compare IC50 values. For graphs that do not fit these criteria, IC50 values and IC50 comparison p values are not listed because these values, while sometimes possible to calculate, are likely inaccurate(58). In these cases, area under curve (AUC) was used to determine if lethal fraction (LF) results were statistically significantly different (GraphPad Prism). AUCs were compared using Welch’s t test (two-tailed), utilizing AUC total area and standard error.

#### Other Statistical Analysis and Software

Statistical analysis and graphing was performed using GraphPad Prism (Version 10.6.1) for all panels except Figure 1A-D, Figure S1A-E, Figure S3A, and Figure 8C,D. Specific statistical tests applied for each dataset are described in the corresponding Figure Legends. Additional detail for Figure 6C, Figure 7C, and Figure S6A: Protein phosphorylation ratios and total protein abundances were log-transformed using the natural logarithm prior to analysis. The primary statistical model was an ordinary least squares (OLS) ANOVA applied to log-transformed ratios, with fixed effects for genotype, dose (treated as a categorical factor with levels 0, 0.5, 1, and 2.5), their interaction, and a blocking factor for gel: log(ratio) ∼ genotype × dose + gel. Pre-specified contrasts compared each genotype pair within every dose level on the log scale. Estimated marginal means and contrasts were obtained using the emmeans package, averaging over gel with equal weights. Inference was performed on the log scale. Because contrasts were planned a priori at specific doses, no multiplicity adjustment was applied. Model assumptions were assessed using Levene’s test across genotype × dose, the Breusch–Pagan test on residuals, and the Shapiro–Wilk test on standardized residuals. To address model heteroscedasticity in the primary analysis, P values were computed using HC3 robust standard errors. When the normality assumption was violated, we performed an additional predefined sensitivity analysis using the Aligned Rank Transform (ART) approach. ART preserves the factorial structure and provides valid inference under non-normality. ART ANOVA and planned contrasts were computed using the *ARTool* package in R(92). Analyses were conducted in R version 4.5.1 (aarch64-apple-darwin20) using the packages *dplyr*, *emmeans*, *car*, *lmtest*, and *sandwich*. BioRender was used to generate Figure 8E.

## Data Availability

All data are contained within the manuscript.

## Acknowledgments

Google Gemini was used exclusively to clarify descriptions of statistical analysis. Simplified Science Publishing.com was used to inform color palette selections for figures. Illustration figures were created with BioRender.com under the institutional license of UMass Chan Medical School and individual academic license of MWL. This work was supported by National Institute of Health (NIH) R01 grants CA229910 and CA290778 (LMS). The content is solely the responsibility of the authors and does not necessarily represent the official views of the National Institutes of Health.

