## Supplemental Figures for "Insulin receptor substrate 2 (IRS2) confers resistance to PI3K pathway inhibition in *PIK3CA* mutant breast cancer"

Figure S1

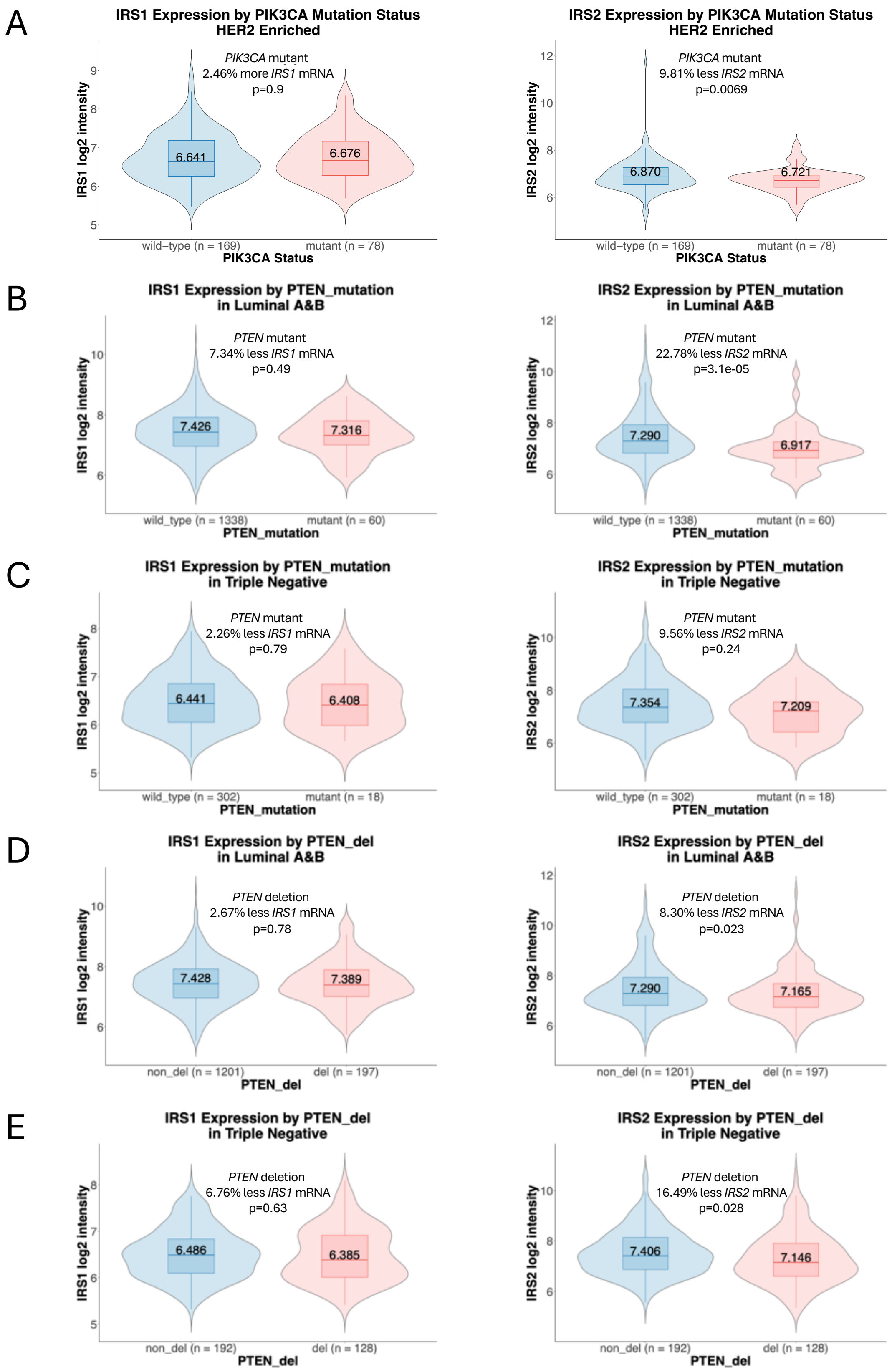
A
IRS1 Expression by PIK3CA Mutation Status
HER2 Enriched
PIK3CA mutant
2.46% more IRS1 mRNA
p=0.9
IRS1 log2 intensity
6.641
6.676
wild−type (n = 169)
mutant (n = 78)
PIK3CA Status
IRS2 Expression by PIK3CA Mutation Status
HER2 Enriched
PIK3CA mutant
9.81% less IRS2 mRNA
p=0.0069
IRS2 log2 intensity
6.870
6.721
wild−type (n = 169)
mutant (n = 78)
PIK3CA Status
B
IRS1 Expression by PTEN_mutation
in Luminal A&B
PTEN mutant
7.34% less IRS1 mRNA
p=0.49
IRS1 log2 intensity
7.426
7.316
wild_type (n = 1338)
mutant (n = 60)
PTEN_mutation
IRS2 Expression by PTEN_mutation
in Luminal A&B
PTEN mutant
22.78% less IRS2 mRNA
p=3.1e-05
IRS2 log2 intensity
7.290
6.917
wild_type (n = 1338)
mutant (n = 60)
PTEN_mutation
C
IRS1 Expression by PTEN_mutation
in Triple Negative
PTEN mutant
2.26% less IRS1 mRNA
p=0.79
IRS1 log2 intensity
6.441
6.408
wild_type (n = 302)
mutant (n = 18)
PTEN_mutation
IRS2 Expression by PTEN_mutation
in Triple Negative
PTEN mutant
9.56% less IRS2 mRNA
p=0.24
IRS2 log2 intensity
7.354
7.209
wild_type (n = 302)
mutant (n = 18)
PTEN_mutation
D
IRS1 Expression by PTEN_del
in Luminal A&B
PTEN deletion
2.67% less IRS1 mRNA
p=0.78
IRS1 log2 intensity
7.428
7.389
non_del (n = 1201)
del (n = 197)
PTEN_del
IRS2 Expression by PTEN_del
in Luminal A&B
PTEN deletion
8.30% less IRS2 mRNA
p=0.023
IRS2 log2 intensity
7.290
7.165
non_del (n = 1201)
del (n = 197)
PTEN_del
E
IRS1 Expression by PTEN_del
in Triple Negative
PTEN deletion
6.76% less IRS1 mRNA
p=0.63
IRS1 log2 intensity
6.486
6.385
non_del (n = 192)
del (n = 128)
PTEN_del
IRS2 Expression by PTEN_del
in Triple Negative
PTEN deletion
16.49% less IRS2 mRNA
p=0.028
IRS2 log2 intensity
7.406
7.146
non_del (n = 192)
del (n = 128)
PTEN_del

**Figure S1. Reduction of *IRS2* mRNA abundance in *PIK3CA* mutant breast cancer.** (A) Differential gene expression analysis of *IRS1* and *IRS2* mRNA in *PIK3CA* WT and *PIK3CA* mutant HER2-enriched breast tumors in the METABRIC dataset. (B and C) Differential gene expression analysis of *IRS1* and *IRS2 mRNA* in *PTEN* WT and *PTEN* mutant luminal A&B (B) and triple-negative (C) breast cancer (METABRIC dataset). (D and E) Differential gene expression analysis of *IRS1* and *IRS2* mRNA in *PTEN* non-deleted (non_del) and *PTEN* deleted (del) luminal A&B (D) and triple-negative (E) breast cancer (METABRIC dataset). Gene expression measurements shown as log2 intensity values (Illumina HT-12 v3 microarray). For all violin plots, the box and whisker plot show the median, 25%(Q1), 75%(Q3) of data (box), Q1 - 1.5 × interquartile range (IQR), and Q3 + 1.5 × IQR of data (whiskers). For all panels, percent changes were determined by comparing the medians of the original data (prior to log transformation). For all panels, statistical analysis by Wilcoxon rank sum test with continuity correction.

Figure S2

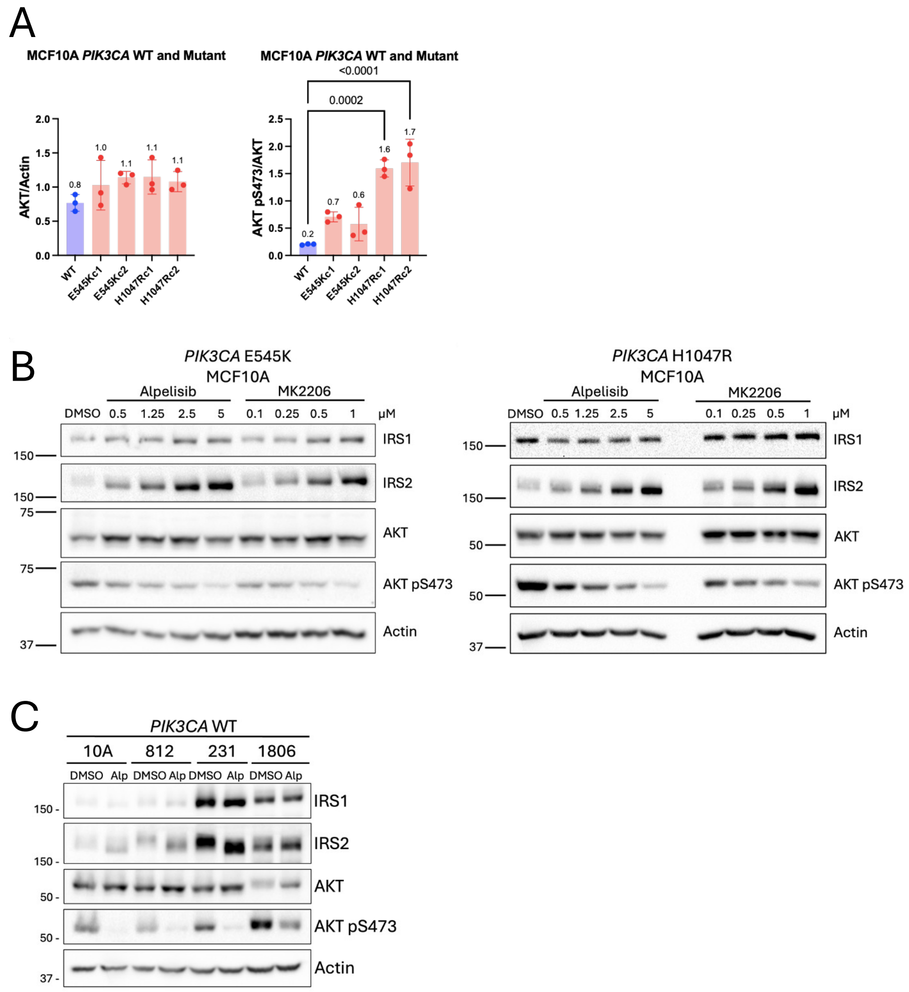


**Figure S2.** ***PIK3CA*** **mutation reversibly suppresses IRS2 mRNA and protein abundance in mammary epithelial cells and breast cancer cell lines.** (A) *PIK3CA* WT and *PIK3CA* mutant knockin MCF10A cells were evaluated for AKT and pS473-AKT expression (see Figure 2B). The data shown represent the mean +/- SD of three independent experiments. Statistical analysis by One-Way ANOVA followed by Dunnett's multiple comparisons test to compare mutants against WT (P values shown). (B) *PIK3CA* E545K and *PIK3CA* H1047R mutant knockin MCF10A cells were treated with increasing concentrations of alpelisib or MK2206 for 24 hrs and cell extracts were evaluated by immunoblot. (C) *PIK3CA* WT breast epithelial and cancer cell lines were treated with or without alpelisib for 24 hrs and cell extracts were evaluated by immunoblot. (10A - MCF10A, 812 - UACC812, 231 – MDA-MB-231, 1806 - HCC1806). Data shown are representative of three independent experiments.

Figure S3

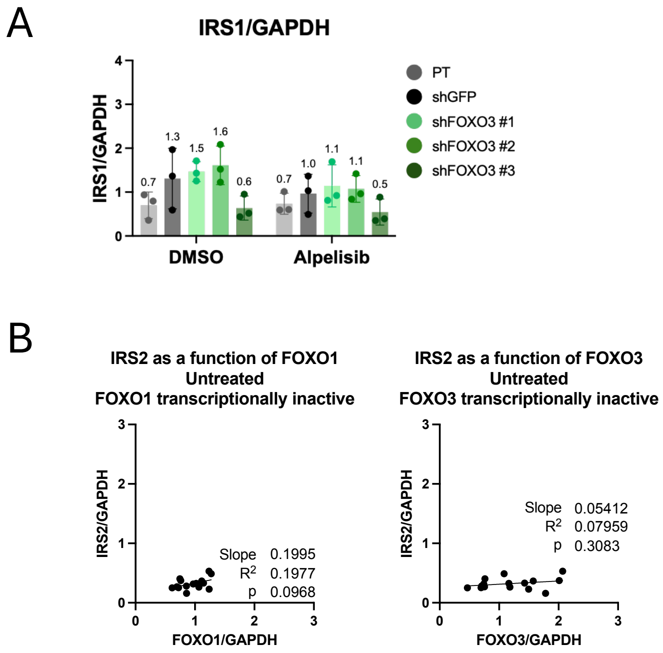


**Figure S3. Mutant PIK3CA regulates IRS2 expression through FOXO3.** (A) *PIK3CA* mutant H1047R MCF10A parental (PT) cells as well as cells expressing control shRNA (shGFP) or shRNA targeting FOXO3 were treated with or without alpelisib (5µM) for 24 hrs and cell extracts were evaluated for IRS1 expression by immunoblot (see Figure 3C). The data shown represent the mean +/- SD of three independent experiments. Statistical analysis by Two-Way ANOVA followed by the Bonferroni test for multiple comparisons, comparing each control (PT and shGFP) to all other sublines within each treatment. (B) Association of IRS2 expression with FOXO1 and FOXO3 abundance in untreated cells (see Figure 3C). Statistical analysis by simple linear regression, $p<0.05$ indicates that the slope is significantly different from zero.

Figure S4

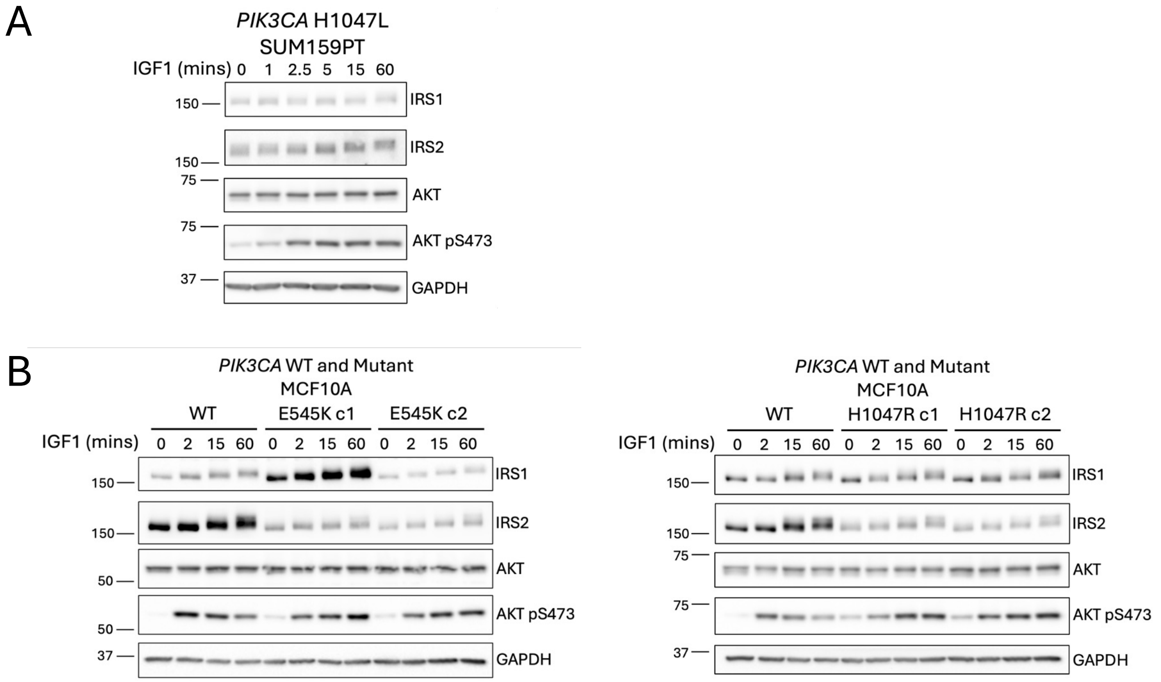


**Figure S4. IGF1 stimulation increases PI3K pathway signaling in *PIK3CA* mutant breast epithelial and cancer cell lines.** (A) SUM159PT cells (*PIK3CA* H1047L) and (B) *PIK3CA* mutant E545K and H1047R MCF10A cells were serum starved overnight, then treated with or without IGF1 (50ng/mL) for increasing amounts of time and cell extracts were evaluated by immunoblot.

Figure S5

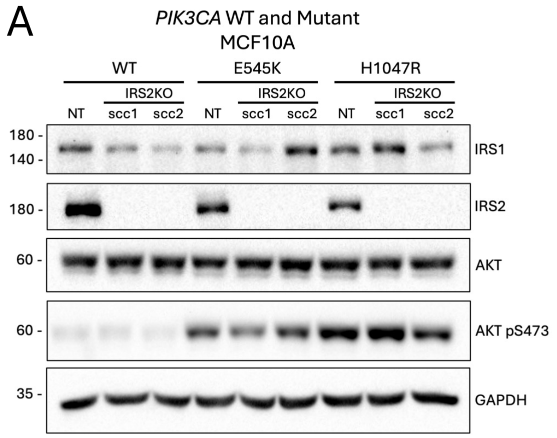


**Figure S5. IRS2 depletion reduces cell proliferation and increases cell death in *PIK3CA* mutant mammary epithelial cells in response to PI3K pathway inhibitors.** (A) MCF10A *PIK3CA* WT and *PIK3CA* mutant cells were treated with non-targeting guide RNA (*NT*) or IRS2-targeting guide RNA (IRS2KO) and cell extracts were analyzed by immunoblot.

Figure S6

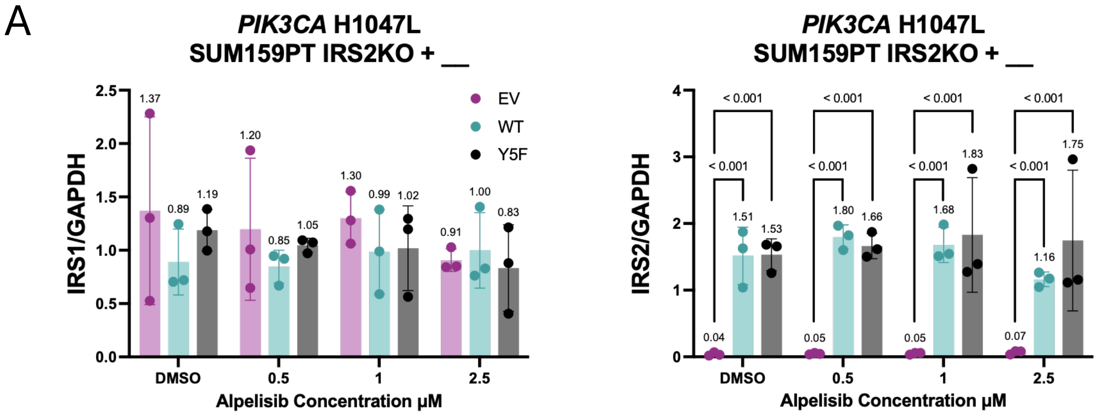


**Figure S6. IRS2-dependent PI3K activation confers resistance to PI3K inhibition in *PIK3CA* mutant breast cancer.** (A) *PIK3CA* H1047L mutant SUM159PT IRS2KO cells expressing empty vector (EV), IRS2-WT or IRS2-Y5F were treated with increasing concentrations of alpelisib for 24 hrs and cell extracts were analyzed for IRS1 and IRS2 expression by immunoblot (see Figure 7C). The data shown in the graphs represent the mean +/- SD of three independent experiments. Statistical analysis was performed using a natural-log transformation followed by ordinary least squares (OLS) ANOVA on log(ratio), with fixed effects for genotype (EV vs. IRS2-WT vs. IRS2-Y5F), dose (categorical: 0 (DMSO), 0.5, 1, 2.5), their interaction, and a gel blocking factor, according to the model: log(ratio) ~ genotype × dose + gel. Pre-specified comparisons between each genotype pair within each dose were conducted on the log scale. Because these comparisons were planned a priori, no multiplicity adjustments were applied (P values shown).

Figure S7

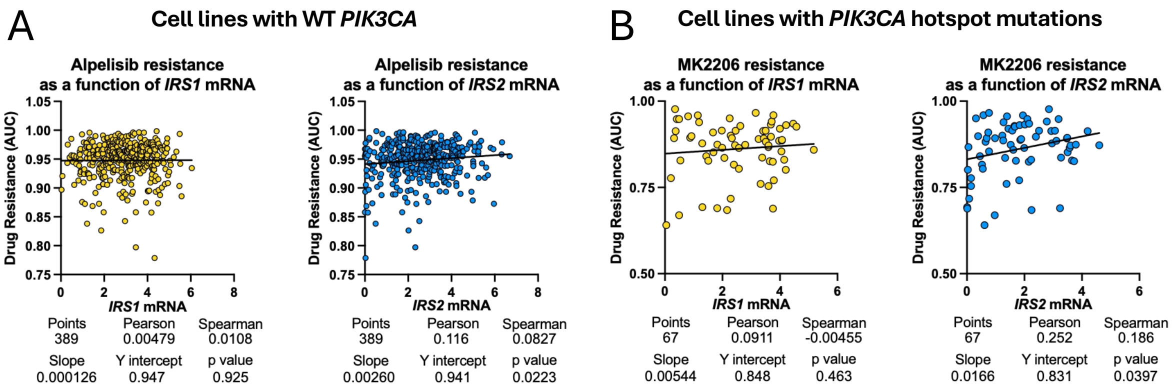


**Figure S7. IRS2 expression positively correlates with resistance to PI3K pathway inhibitors in *PIK3CA* mutant cancer cell lines.** (A) Correlation between *IRS1* and *IRS2* mRNA and sensitivity to alpelisib in cancer cell lines with WT *PIK3CA* (excluding myeloid and lymphoid-derived cell lines) in the DepMap dependency dataset. (B) Correlation between *IRS1* and *IRS2* mRNA and sensitivity to MK2206 in cancer cell lines with *PIK3CA* hotspot mutations (excluding myeloid and lymphoid-derived cell lines) in the DepMap dependency dataset. For (A) and (B), P values shown are from simple linear regression, $p<0.05$ indicates that the slope is significantly different from zero. mRNA abundance shown is log2(TPM+1).
